# Bidirectional disruption of Lrrk2 function drives T cell dysregulation and an exhaustion-like immune response

**DOI:** 10.64898/2026.08.16.745139

**Authors:** Robert C. Sharp, Shannon C. Wall, Jordan C. Follet, Isaac B. Deng, Matthew J. Farrer

## Abstract

**Background:** Neurodegenerative diseases including Parkinson’s disease (PD) are increasingly associated with dysfunction in both central and peripheral immune systems. Pathogenic mutations in leucine-rich repeat kinase 2 (*LRRK2*) represent a major cause of familial PD, while common polymorphisms are associated with inflammatory diseases.

**Methods:** Here, using immunophenotyping flow cytometry and quantitative PCR (qPCR), we compared immune cell populations and function across both the central and peripheral immune systems in C57BL/6J wild type (WT), *Lrrk2* p.G2019S knock-in (GKI) and *Lrrk2* knock-out (LKO) models in basal and *ex vivo* immune-stimulated conditions.

**Results:** With a focus on T cell biology, compared to their WT counterparts at baseline, GKI mice exhibit higher populations of Cd8^+^ and T_H17_ T cell subsets in the brain, whereas LKO mice exhibited unique central memory (T_CM_), follicular helper (T_FH_), and T_H2_ lineages. In the periphery, GKI mice demonstrated higher T_H1_, T_H2_, and T_H17_ subset expansions, whereas peripheral alterations in LKO are largely restricted to T_H17_ subsets. Within mutant genotypes, a striking discrepancy was observed between baseline gene expression and the translated proteins encoded, that reveals a fundamental loss of basal immune homeostasis. This phenomenon was further exposed following an acute (6-hour) *ex vivo* lipopolysaccharide (LPS) immune challenge. Following stimulation, GKI immune cells had reduced transcription, alongside stalled translation, for almost all effector molecules examined, while LKO immune cells had fewer transcriptional changes compared to wild type. Overall, both mutant lines had stalled or flatline effector molecule production after immune stimulation, suggesting a profound loss of functional responsiveness. This hypothesis was supported by a significant increase in surface protein of the inhibitory receptor Pd-1 on regulatory T cells (T_REG_) and T_H1_/T_H2_ Cd4^+^ T cell subsets in GKI mice. LKO immune landscapes trended toward similar exhaustion patterns, albeit less evident.

**Conclusions:** These data suggest that bidirectional disruptions to normal Lrrk2 function break immune homeostasis. Immune cell function should be carefully considered when targeting LRRK2 kinase activity in patients with PD.

## BACKGROUND

In neurodegenerative diseases, neurologic pathologies are associated with local and systemic immune responses. In Parkinson’s disease (PD), both the midbrain and peripheral immune systems also utilize dopamine signaling to dynamically regulate physiologic responses within and between these tissues[1–3]. Dysregulation of this dopaminergic-immune crosstalk is increasingly recognized as a key driver of neuroinflammation[4]. However, while the loss of dopaminergic signaling undoubtedly impacts immune behavior indirectly, the direct, cell-intrinsic role of PD-linked mutations on immune cell functionality remains a critical factor.

Pathogenic mutations in leucine-rich repeat kinase 2 (*LRRK2*) are enriched across distinct human populations[5–7] and is highly expressed in myeloid cells and modestly in lymphoid cells[8–13]. The geographic distribution of the *LRRK2 c.6055G>A* (encoding p.G2019S) variant is consistent with antagonistic pleiotropy – the concept that positive selection enhances early survival[14] although here the late-life cost is increased risk for PD[5–7]. It remains unclear how precisely LRRK2 activity impacts immune subset differentiation or the production of specific effector molecules including interleukins. However, chronic LRRK2 kinase activation may drive cellular senescence[15–17] and hamper appropriate immune responses.

In this study, we compared immune cell presence, memory differentiation, and effector molecule production in C57BL/6J wild type (WT), *Lrrk2 c.6055G>A* (GKI)[18] and *Lrrk2* knock out (LKO) mice[19]. We hypothesized that the *Lrrk2* p.G2019S kinase-activating mutation and the complete loss of *Lrrk2* will uniquely disrupt immunoregulatory pathways, resulting in distinct alterations to central and peripheral immune compositions and effector molecule production. Experiments were performed using mice subject to standard vivarium housing conditions (in microisolator caging with forced HEPA-filtered air, sterilized bedding and standard chow) in a basal state. We used quantitative polymerase chain reaction (qPCR) and flow cytometry immunophenotyping to assess gene and protein expression of immune cell subset markers and effector molecules from whole brain and peripheral blood/splenocyte-derived populations. Our findings show Lrrk2 impacts the regulation of central and peripheral immune systems, preferentially targeting T cell biology.

## METHODS

### Mouse Models

The following strains were used for this study: Wild-Type (WT) C57BL/6J, C57BL/6J-B6.Cg-*Lrrk2^tm1.1Hlme^*/J (GKI mouse, strain # 030961)[18], and C57BL/6-*Lrrk2^tm1Mjfa^*/J (LKO mouse, strain #012444)[19]. All mouse models are available from Jackson Laboratory©. GKI and LKO mice have been maintained on the same C57BL/6J mouse background for over 10 generations, where studies were approved by the Institutional Animal Care and Use Committee (IACUC) at the University of Florida. Following the National Institute of Health (NIH) guidelines, all mice were kept on a reverse light cycle and housed in single-sex group-housed in cages after weaning. For all experiments, male mice aged 3-4 months were used for each experimental assay (N = 5 WT; 5 GKI; 5 LKO for each experimental assay, totaling 45 mice). Each mouse provided brain and peripheral blood/splenocyte immune cells for each assay. Genotype validation was completed via ear notch DNA extraction as previously described[18, 19].

### Brain, Peripheral Blood, and Spleen Isolation

Mice were humanely euthanized by Isopire™ (isoflurane; Dechra®) and transcardially perfused with cold Hank’s Balanced Salt Solution (HBSS; Gibco™) with 5% fetal bovine serum (FBS; Gibco™) at a rate of 10 mL/min (up to 30 mL). Exsanguinated diluted peripheral blood was collected in a 50 mL conical tube and the spleens were removed and placed within the same tube. Whole brain extractions were conducted in a 50 mL conical tube containing cold HBSS with 5% FBS.

### Brain Immune Cell Isolation

Extracted brains were placed in cold HBSS with 5% FBS and transferred into a large cell culture dish, where they were sliced with a scalpel into approximately ten different slices. Slices in cold HBSS with 5% FBS were mechanically dissociated and filtered through a 100 μm cell strainer (Corning Life Sciences©) while being washed with additional cold HBSS with 5% FBS. Single-cell suspensions were then centrifuged for 300g for 6 mins at 4°C and cell pellets were resuspended in Accumax™ Cell Dissociation Solution (Innovative Cell Technologies®) and incubated at room temperature on a rocker for 30 mins. Following incubation, the single-cell suspensions were filtered again through a 70 μm cell strainer (Corning Life Sciences©) and washed with cold HBSS with 5% FBS. Single-cell suspensions were then centrifuged once more at 300g for 6 mins at 4°C. Next, the cell pellet was mixed with a 30% isotonic Percoll® solution (MP Biomedicals Inc©). Single-cell suspensions with the 30% isotonic Percoll® solution were then centrifuged with slow acceleration and no brake at 500g for 30 mins at room temperature. After centrifugation, the top myelin layer was removed and the Percoll® suspended immune cell, glial, and neuron cell pellet were washed with homogenized cell pellets in cold HBSS with 5% FBS by repeated centrifugation steps at 250g for 6 mins a 4°C.

### Peripheral Blood Processing/Spleen Homogenization and Immune Cell Isolation

Peripheral blood/spleen were mechanically homogenized, filtered through a 100 μm cell strainer, and washed with HBSS with 5% FBS. The single-cell suspensions were centrifuged using a slow acceleration and no brake at 500g for 30 mins at 4°C, where the supernatant was then removed. The resulting immune cell/splenocyte pellets were resuspended in HBSS with 5% FBS and treated with Ficoll®-Paque Premium 1.084 gradient solution (Millipore Sigma©). The single-cell suspensions with Ficoll® were centrifuged using a slow acceleration and no brake at 500g for 30 mins at room temperature. Isolated peripheral blood mononuclear cells (PBMCs) were collected from the middle buffy coat layer of the gradient and washed with HBSS with 5% FBS after repeated centrifugation steps at 250g for 6 mins at 4°C.

### RNA Extraction, cDNA Synthesis, and qPCR

RNA extractions and complementary DNA (cDNA) synthesis was performed on both brain and peripheral single-cell suspensions as previously reported[20]. Briefly, single-cell suspensions were homogenized using QIAshredder (Qiagen©) tubes and RNA was isolated following the protocols provided by the RNeasy® Mini Kit (Qiagen©). Purified RNA was quantified using a NanoDrop™ One/One^C^ spectrophotometer (Thermo Fisher Scientific©). A total of 400 ng of RNA per sample was converted into cDNA using protocols from the High-Capacity cDNA Reverse Transcription Kit (Applied Biosystems™). For qPCR, 4 μL of cDNA from each sample was pipetted into 384-well plates in triplicates for each gene examined with the following master mix solution: 10 μL of TaqMan™ Universal Master Mix II, with UNG (Applied Biosystems™), 5 μL of RNase/DNase-free water, and 1 μL of TaqMan™ Gene Expression Assay (Thermo Fisher Scientific©). TaqMan™ Gene Expression Assay IDs used in this study can be found in **Supplementary Table 1**. Quantification of each sample were conducted and analyzed on a QuantStudio™ 5 Real-Time PCR System (Applied Biosystems™) using QuantStudio™ Design & Analysis Software (Applied Biosystems™). Comparative Ct values (ΔΔCt) for each sample were calculated by comparing ΔCt (Ct value of targeted gene – Ct value of combined means of *Gapdh* and *Actb* [tested for significance between each group using a Welch’s ANOVA with Dunnett’s T3 post-hoc test]) across all samples to the WT group. The formula used is: ΔΔCt = ΔCt of targeted gene from experimental group – ΔCt of targeted gene from control group. Relative gene expression for each gene of interest from each sample was then calculated using the 2^(−ΔΔCt)^ formula.

### Staining Cells for Flow Cytometry

Brain and PBMC single-cell suspensions were transferred into separate 5 mL Falcon® round-bottom polystyrene flow cytometric tubes (Corning©) in duplicates. The samples were washed with PBS and centrifuged for 5 mins at 4°C at 350g. Supernatants were discarded and 0.2 μL of Live/Dead™ Fixable Near-IR Dead Cell Statin (Invitrogen™) was added to each cell pellet in 50 μL of PBS. The cells were then incubated with Live/Dead stain for 15 mins at 4°C, washed with flow cytometry stain buffer (1X PBS, 2%FBS, and 0.05% NaN_3_), and centrifuged. Following supernatant removal, cell pellets were treated with 0.1 μL of TruStain FcX™ Plus solution (anti-mouse Cd16/32; Biolegend®) in 100 μL of stain buffer and incubated for 10 mins at 4°C. Extracellular flow cytometry antibodies (**Supplementary Table 2**) were then added at 1 μL per marker to each sample tube. Unstained controls and fluorescent minus one (FMO) control tubes were created for compensation and gating purposes. After adding the antibody mixtures, the samples were incubated for 45 mins at 4°C. Samples were then washed with 1 mL of stain buffer and centrifuged at 350g for 5 mins at 4°C. Supernatants were discarded and the cell pellets were processed using the True-Nuclear™ Buffer Set (Biolegend®). Samples were treated with 250 μL of 1X fixative buffer for 20 mins at room temperature. After incubation, 1 mL of 1X permeabilization buffer was then added and the samples were then centrifuged for 400g for 5 mins at room temperature. Supernatants were then discarded, and cell pellets were resuspended in 250 μL of 1X permeabilization buffer and incubated for 20 mins at room temperature. Samples were then centrifuged at 400g for 5 mins at room temperature and supernatants were discarded. The cell pellets were then treated with 100 μL of 1X permeabilization buffer and 1 μL of each of the intracellular flow cytometry antibodies per marker are added (**Supplementary Table 2**). The samples were then incubated in the antibody mixtures for 45 mins at room temperature. After incubation, 1 mL of 1X permeabilization buffer was added to the samples and were centrifuged at 450g for 5 mins at room temperature. The supernatants were then discarded, and the wash step was repeated twice. Once fully washed, the cell pellets were resuspended in 300 μL of stain buffer and were incubated overnight at 4°C. The following day, single-color controls using both UltraComp eBeads™ Plus Comp Beads (Invitrogen™) for flow antibodies and ArC™ Amine Reactive Comp Beads (Invitrogen™) for Live/Dead staining were prepared for compensation purposes. Single-color controls, unstained controls, FMOs, and the samples underwent flow cytometry analysis on the Cytek® Aurora 5 laser [16UV-16V-14B-10YG-8R].

### Acute *Ex vivo* Lipopolysaccharide Treatment of Isolated Immune Cells for Intracellular Effector Molecule Production

Brain and peripheral single cell suspensions underwent acute *ex vivo* stimulation for 6-hours to initiate the production of effector molecules, which are otherwise difficult to detect by flow cytometry at baseline[21, 22]. The samples were resuspended in complete Dulbecco’s Modified Eagle Medium/Nutrient Mixture F-12 (cDMEM/F-12), which consisted of a DMEM/F-12 base and contained the following reagents: 10% FBS, 1X GlutaMAX™ (Gibco™), 1X Penicillin/Streptomycin (Corining®), 1 mM Sodium Pyruvate (Gibco™), 1X Non-Essential Amino Acids (Gibco™), 1 mM of HEPES (Gibco™), 2 μL of 2-mercaptoethanol (BME), and 1.2 mL of Sodium Hydroxide Solution (Honeywell International©). Samples were plated in duplicates on 12-well or 24-well plates (depending on cell number) and were treated with 1X lipopolysaccharide (LPS) solution/mL (2.5 μg LPS/mL; Invitrogen™) and 0.65 μL/mL of GolgiStop™ Protein Transport Inhibitor (BD Biosciences©) for 6-hours at 37°C with 5% CO_2_. Unstimulated controls without LPS treatment were also conducted to determine proper flow cytometry gating. Following incubation, cell suspensions were removed from the plate, where the macrophage detachment solution (PromoCell®) was used according to manufacturer’s instructions to remove any adherent cells from the plates. Previously adherent cells were then combined with the cell suspensions and samples were then centrifuged for 350g for 5 mins at 4°C. The supernatants were discarded, and the cell pellets were washed twice with PBS. The samples were then treated for flow cytometry staining and analysis as previously described above using a specialized effector panel (**Supplementary Table 3**).

### Flow Cytometry Data Processing

Flow cytometry standard (FCS) files were processed using the FlowJo™ software package (BD Life Sciences©, v10.10.0). All gating for each flow cytometry panel and subsequent analysis was based upon unstained controls, unstimulated controls (for the intracellular effector molecule production panel), single color controls, and FMOs. Automated compensation for each flow panel was performed using the results of the controls within the FlowJo™ software. Examples of gating strategies (created from concatenated FCS files of all samples) for detection of median fluorescent intensity (MFI) and immune subset cell counts can be found in the following figures: MFI readings for immune subset markers in brain single-cell suspension (**Supplementary Figure 1**); MFI readings for immune subset markers in peripheral single-cell suspension (**Supplementary Figure 2**); immune subset cell counts in brain single-cell suspension (**Supplementary Figure 3**); immune subset cell counts in peripheral single-cell suspension (**Supplementary Figure 4**); MFI readings for intracellular effector molecule production in brain single-cell suspension (**Supplementary Figure 5**); MFI readings for intracellular effector molecule production in peripheral single-cell suspension (**Supplementary Figure 6**). To compare relative protein expression across genotypes, MFI readings were normalized to the background MFI derived from the unstained/unstimulated controls (nMFI = sample MFI - unstained/unstimulated control MFI). For comparisons of immune cell subset populations (complete list of phenotypes and references used for characterization found in **Supplementary Table 4**), population percentages were calculated from total live single cells in each. Integrated MFI (iMFI) was calculated to reflect both marker intensity and population frequency using the equation: iMFI = (nMFI value of maker) * (percentage positive for marker within that subset)[23, 24]. To the best of our knowledge, we have reported all necessary methods and panel optimizations according to the Minimum Information about a Flow Cytometry Experiment (MIFlowCyt) protocol[25, 26].

### T-distributed Stochastic Neighbor Embedding (tSNE) Clustering

tSNE clustering for the immune subset flow cytometry panel (**Supplementary Table 2**) was performed using concatenated (combined) FCS files in FlowJo™. Workflow for tSNE cluster identification for both brain and peripheral single-cell suspensions can be found in **Supplementary Figure 7**. Briefly, the concatenated samples were reduced to 1000000 events (using DownSample plugin, BD Life Sciences©, V3), gated on live single cells, and subjected tSNE clustering. The X-Shift plugin (BD Life Sciences©, v1.4.1) was used to determine the meta-cluster numbers from each tSNE plot and FlowSOM (BD Life Sciences©, v3.0.18) was utilized to categorize the meta-clusters. Clusters were then identified using Cluster Explorer (available in FlowJo™), where clusters were classified based on relative expression levels of each immune marker examined, where ≥10^4^ relative expression level cutoff was chosen based upon FMO and unstained control relative expression levels. Clusters were named first by one of the following “backbone” immune cell subset marker: Cd8^+^ (Cd8^+^ T cells), Cd4^+^ (Cd4^+^ T cells), Cd11c^+^ (DCs), Nkp46^+^ (NK cells), Cd22^+^ (B cells), Cd11b^+^ (Cd11b^+^ phagocytes), Tmem119^+^ (Tmem119^+^ phagocytes), Iba1^+^ (Iba1^+^ phagocytes), Gfap^+^ (Gfap^+^ astrocytes), and S100b^+^ (S100b^+^ astrocytes). Once each cluster was named based on a “backbone” immune cell subset marker, further identification of any additional markers within each cluster that had ≥10^4^ relative expression levels were labeled for each cluster. From each fully labeled cluster, WT, GKI and LKO cells were identified within the tSNE plots and were compared to the normalized cluster population (expressed as a fraction of total percentage) for each cluster (**Supplementary Figure 7**).

### Statistical Analysis

Graphing and statistical analysis were performed using GraphPad Prism© (v10.5.0) software. For parametric data, statistical analysis was done using a Brown-Forsythe and Welch ANOVA with a Dunnett’s T3 multiple comparisons test. For nonparametric data, a Kruskal-Wallis test with a Dunn’s multiple comparison test was utilized. A Benjamini-Hochberg FDR correction was applied for all *p-values* across all transcriptional and relative protein expression tests.

## RESULTS

### Distinct Profiling of Central and Peripheral Immune Systems Reveal Basal T cell Remodeling Across *Lrrk2* Genotypes

We initially evaluated mononuclear immunophenotyping markers in an exploratory, unbiased manner to assess central (brain) and peripheral immunity (PBMCs/splenocytes) in GKI and LKO compared to WT controls under basal conditions. We conducted qPCR for relative gene expression analysis and flow cytometry immunophenotyping of individual immune cell subset markers on live single cells before separating them by cellular subset.

Using the 2^(−ΔΔCt)^ method for qPCR gene expression analysis (**Figure 1**), we targeted a panel of core “backbone” immune cell markers (**Supplementary Table 1**) including: *Cd3e* (Cd3 epsilon; T cell receptor), *Cd8a* (Cd8 alpha; Cd8^+^ T cells), *Cd4* (Cd4^+^ T cells), *Itgax* (Integrin alpha x or Cd11c; Cd11c^+^ dendritic cells [DCs]), *Ncr1* (Natural cytotoxicity triggering receptor 1 or Nkp46; Nkp46^+^ natural killer [NK] cells), *Cd22* (B cells), *Itgam* (Integrin alpha m or Cd11b; Cd11b^+^ monocytes/macrophages/microglia), *Tmem119* (Transmembrane protein 119; Tmem119^+^ phagocytes), *Aif1*, (Allograft inflammatory factor 1 or ionized calcium binding adapter molecule 1 [Iba1]; Iba1^+^ phagocytes), *Gfap* (Glial fibrillary acidic protein; Gfap^+^ astrocytes), and *S100b* (S100 calcium-binding protein b; S100b^+^ astrocytes). Among these, *Cd3e*, *Cd8a*, and *Cd4* were significantly altered in both the brain (**Figure 1A**) and periphery (**Figure 1B**) of GKI and LKO mice. Interestingly, within the GKI mice, all three genes were significantly upregulated in the brains (**Figure 1A**). However, the same genes were significantly downregulated within the periphery (**Figure 1B**). In contrast, LKO mice demonstrated significant upregulation of *Cd8a* in brain (**Figure 1A**), while *Cd3e*, *Cd8a*, and *Cd4* expression were concurrently decreased in the periphery (**Figure 1B**). Notably, almost all targeted immune markers examined presented significantly lower relative gene expression in the periphery of GKI mice (**Figure 1B**).

**Figure 1:**
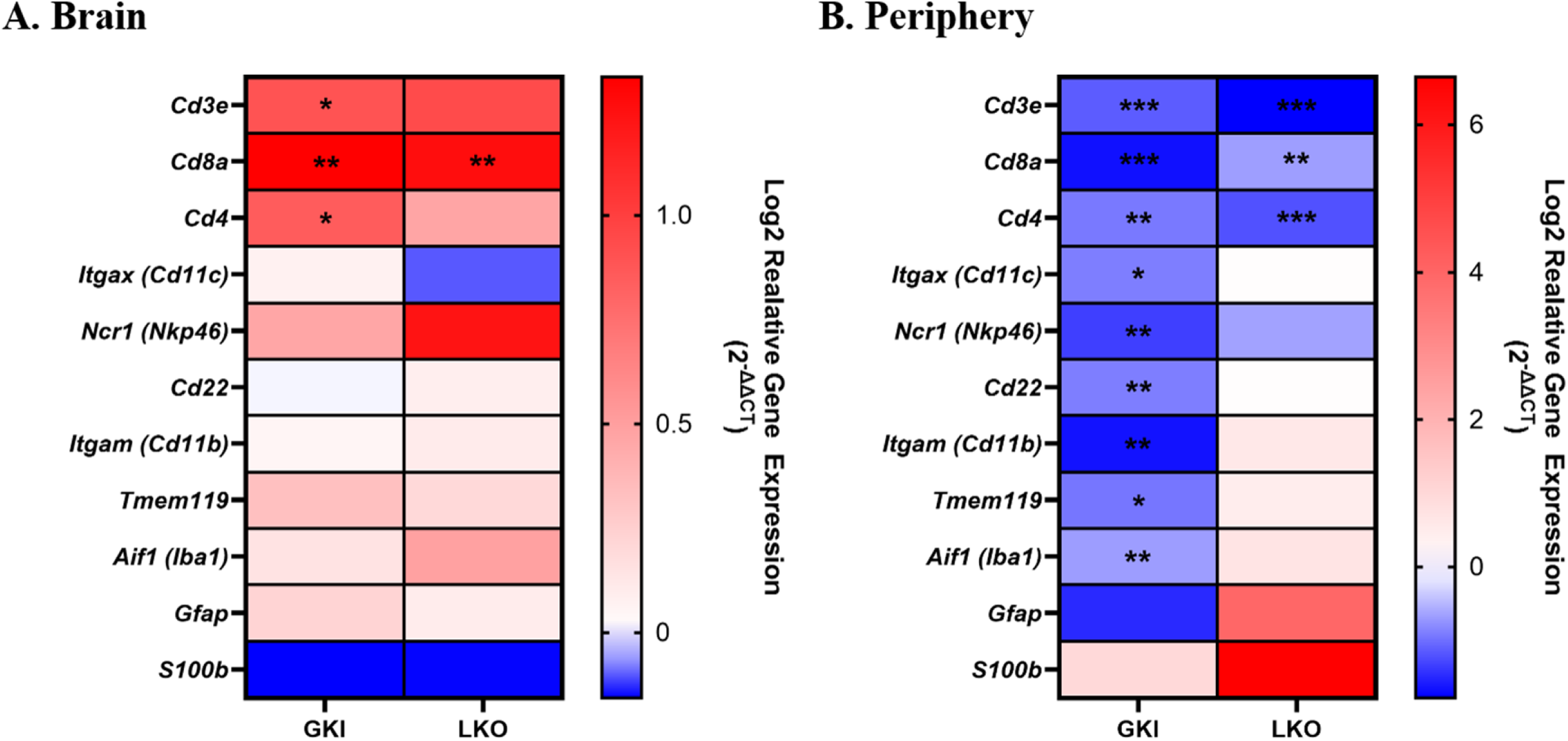
Divergent Transcriptional Profiles in Central and Peripheral Immune Systems Reveal Baseline Immune Discrepancies Across *Lrrk2* Genotypes. (**A.**) Central (whole brain) and (**B.**) peripheral (PBMCs/splenocytes) gene expression profiles of lineage and glial markers under basal conditions (N = 5 per genotype). Data are presented as heatmaps of Log2 of relative gene expression (2^−ΔΔCT^) normalized to WT controls. Statistical analysis was done using a Brown-Forsythe and Welch ANOVA with a Dunnett’s T3 multiple comparisons test. A Benjamini-Hochberg FDR correction was applied for all *p-values* across all comparisons. Significance symbols: *: *P-value ≤ 0.05*; **: *P-value ≤ 0.01*; ***: *P-value ≤ 0.001*

Spectral flow cytometry was performed to assess the relative protein expression (described as nMFI values) of our complete immunophenotyping panel (**Supplemental Table 2**) in live single cells before categorizing the cells into immune subsets (**Figure 2**, see **Supplementary Figure 1** and **Supplementary Figure 2** for gating strategies). Evaluating the Log2 fold change of nMFI values (Log2 nMFI FC) between mutant genotypes and WT controls demonstrated the most pronounced shifts in protein expression occurred in markers associated with T cells. In GKI brains, we found significantly higher protein fold changes for Cd3^+^, Cd8a^+^, Cd183^+^ (Cxcr3^+^, T helper 1 [T_H1_] T cell associated), and Cd62l^+^ (L-selectin, memory T cell associated) (**Figure 2A**). LKO brains exhibited no significant protein fold changes (**Figure 2A**). In the periphery (**Figure 2B**), GKI mice displayed significant protein fold changes for Cd196^+^ (Ccr6^+^, T helper 17 [T_H17_] associated) and Pd-1^+^ (immune cell inhibition and T follicular helper [T_FH_] associated). Peripheral LKO profiles demonstrated significant increases in overall Cd3^+^ (T cell receptor, bulk T cell marker) and Pd-1^+^ protein fold change, but a significant decrease in Cd185^+^ (Cxcr5^+^, T_FH_ associated) (**Figure 2B**).

**Figure 2:**
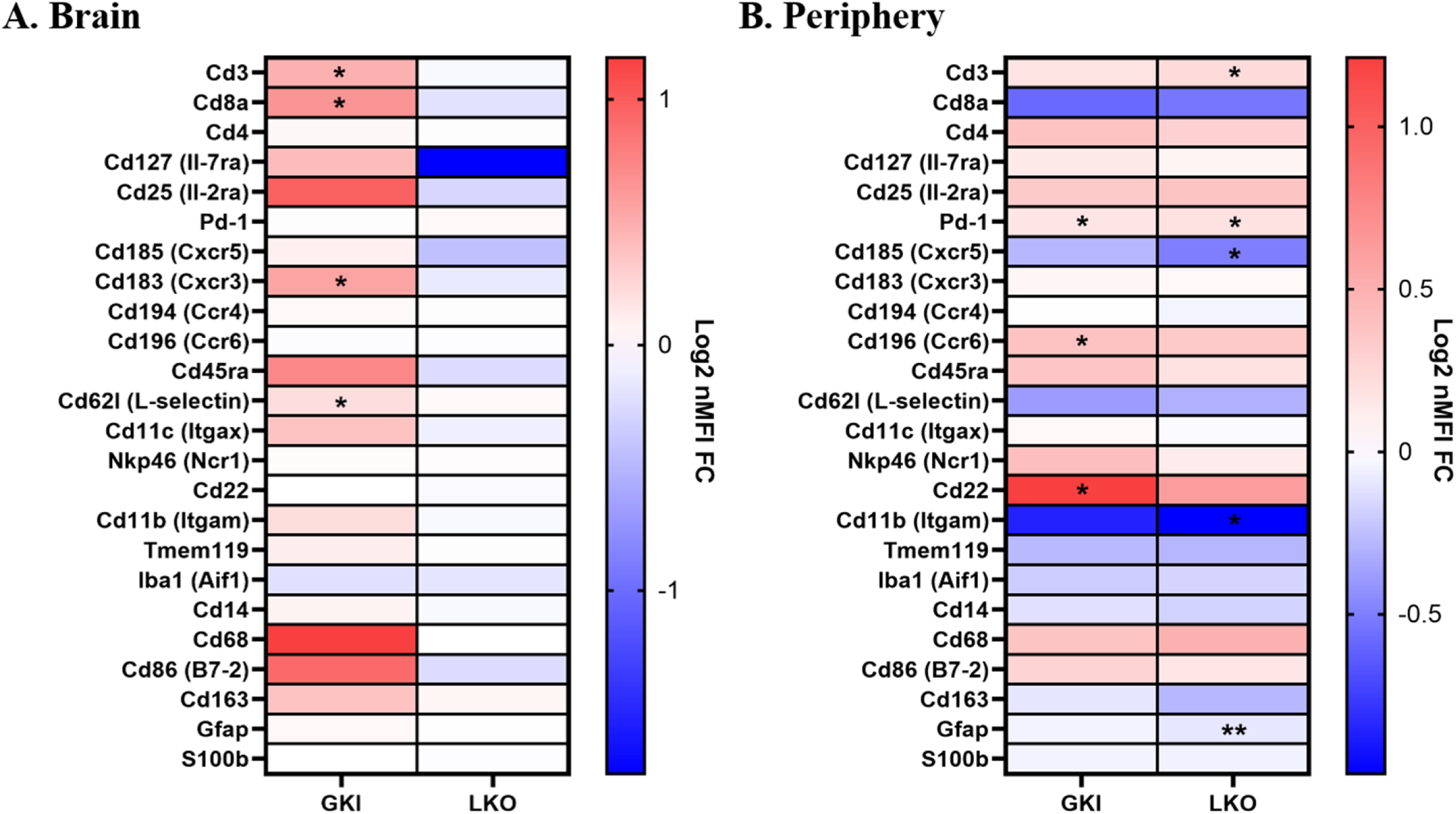
Baseline T cell Surface Protein Profiles in Central and Peripheral Immune Systems Are Altered Across *Lrrk2* Genotypes. (**A.**) Central (whole brain) and (**B.**) peripheral (PBMCs/splenocytes) relative protein expression profiles across a 24-marker spectral flow cytometry panel in overall live single cells (N = 5 per genotype). Heatmaps display the mean Log2 normalized median fluorescent intensity fold change (Log2 nMFI FC) relative to WT controls. For parametric data, statistical analysis was done using a Brown-Forsythe and Welch ANOVA with a Dunnett’s T3 multiple comparisons test. For nonparametric data, a Kruskal-Wallis test with a Dunn’s multiple comparison test was utilized. A Benjamini-Hochberg FDR correction was applied for all *p-values* across all comparisons. Significance symbols: *: *P-value ≤ 0.05*; **: *P-value ≤ 0.01*

We next implemented an unbiased tSNE clustering analysis via FlowSOM/X-shift to confirm unique immune cell clustering (**Figure 3**; **Supplementary Figure 7** for workflow). In GKI brains, unique Cd8^+^ T cell grouping (Cluster 8 and Cluster 10) emerged relative to WT cells (**Figure 3A; Supplementary Table 5** for detailed statistics). The phenotypes identified from these clusters suggest a prolonged proinflammatory response, marked by elevated levels of traditional T cell markers as Cd25^+^ (Il-2ra), Cd45ra^+^, Cd183^+^ (Cxcr3^+^) and Cd127^+^ (Il-7ra) and non-traditional T cell co-stimulatory markers such as Cd86^+^ (B7-2^+^)[27–29] and Cd11c^+^ (Itgax^+^)[30–32] (**Figure 3A**; **Supplementary Table 5** for detailed statistics). Notably, GKI brains also featured a significant decrease in a rarer immune population, which is a double-positive (Cd4^+^Cd8^+^) T cell cluster (Cluster 12)[33–35]. Conversely, peripheral tSNE clustering analysis for LKO mice displayed significant expansions of unique Cd8^+^ effector (Cd45ra^+^ Cd68^+/−^) T cell clustering (Cluster 7 and Cluster 8) and in Pd-1^+^ Cd4^+^ T cell clustering (Cluster 12; **Figure 3B**; **Supplementary Table 5** for detailed statistics).

**Figure 3:**
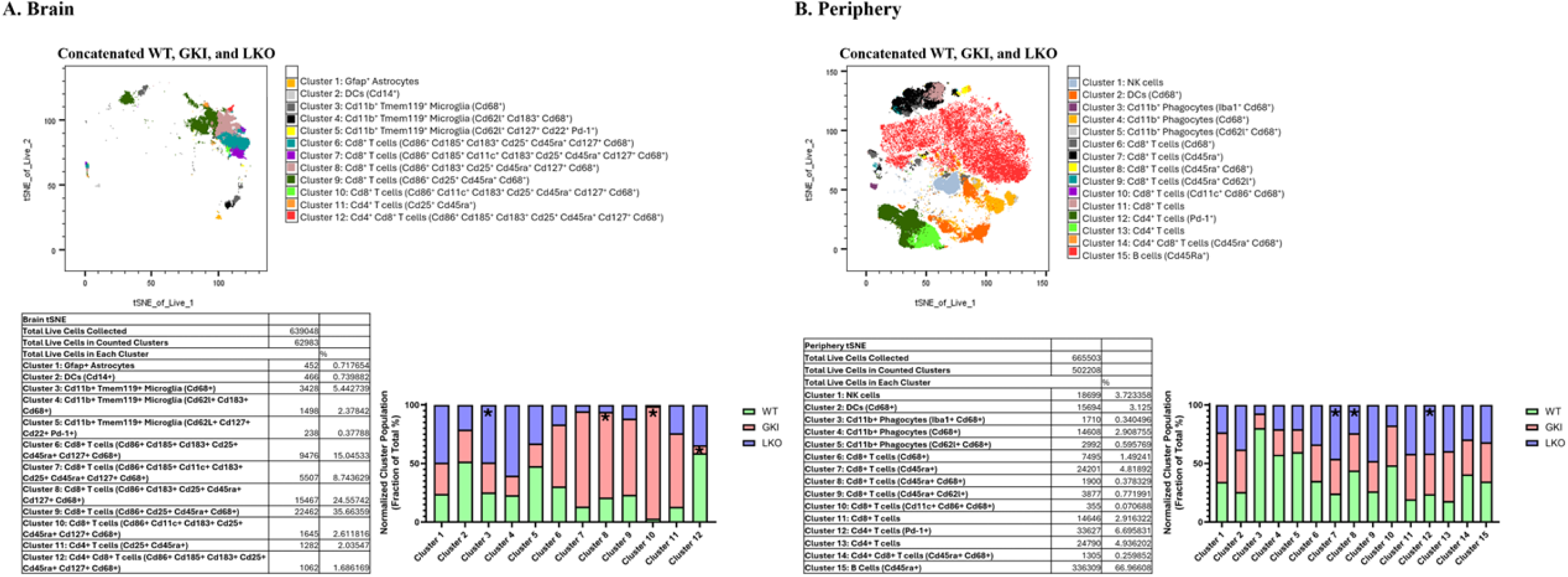
Unbiased tSNE Topologies Capture Distinct *Lrrk2* Genotype-Specific T cell Cluster Landscapes at Baseline Central and Peripheral Immune Systems. High-dimensional tSNE clustering of concatenated spectral flow cytometry datasets from (**A.**) brain (639,048 total live cells collected across cohorts) and (**B.**) peripheral (665,503 total live cells collected across cohorts) mononuclear cell populations (N = 5 per genotype). Topographic maps display phenotypic distribution across 12 different algorithmically identified central meta-clusters and 15 peripheral meta-clusters via FlowSOM clustering. Accompanying quantification tables detail exact cell counts and total percentages per cluster. Bottom bar graphs illustrate the relative fraction of total population for each shifting cluster across WT, GKI, and LKO genotypes, highlighting significant visual topography shifts in mutant mice. For parametric data, statistical analysis was done using a Brown-Forsythe and Welch ANOVA with a Dunnett’s T3 multiple comparisons test. For nonparametric data, a Kruskal-Wallis test with a Dunn’s multiple comparison test was utilized. Significance symbols: *: *P-value ≤ 0.05*

Results of gene (qPCR) and protein (flow cytometry) expression indicate that, without immune stimulation, markers identifying T cells are most significantly altered in the central and peripheral immune systems of GKI and LKO mice compared to WT. Hence, further investigation focused on T cell biology.

### *Lrrk2* Genotypes Drive Divergent T cell Subset Accumulations Within the Brain

Using immune cell subset targeted flow gating (**Supplementary Figure 3**), we quantified distinct immune subsets within the pool of total live brain cells to determine the exact nature of T cell accumulation (see **Supplementary Figure 8A** for analysis of all immune cell subsets) using a defined immunophenotyping panel (**Supplementary Table 4**). Overall, GKI brains exhibited significantly more T cell populations, which were predominantly Cd8^+^ (**Figure 4A**; **Supplementary Figure 9** for flow gating of significant findings). Notably, most Cd8^+^ T cells along with T_H17_ T cells expressed a more naïve-like (Cd45ra^+^Cd62l^+^) phenotype (**Figure 4A**). LKO brains showed completely distinct T cell expansions, where significant increases in bulk central memory (T_CM_) T cells, specifically T_FH_ subsets, and overall T helper 2 (T_H2_) cell frequencies occurred (**Figure 4B**).

**Figure 4:**
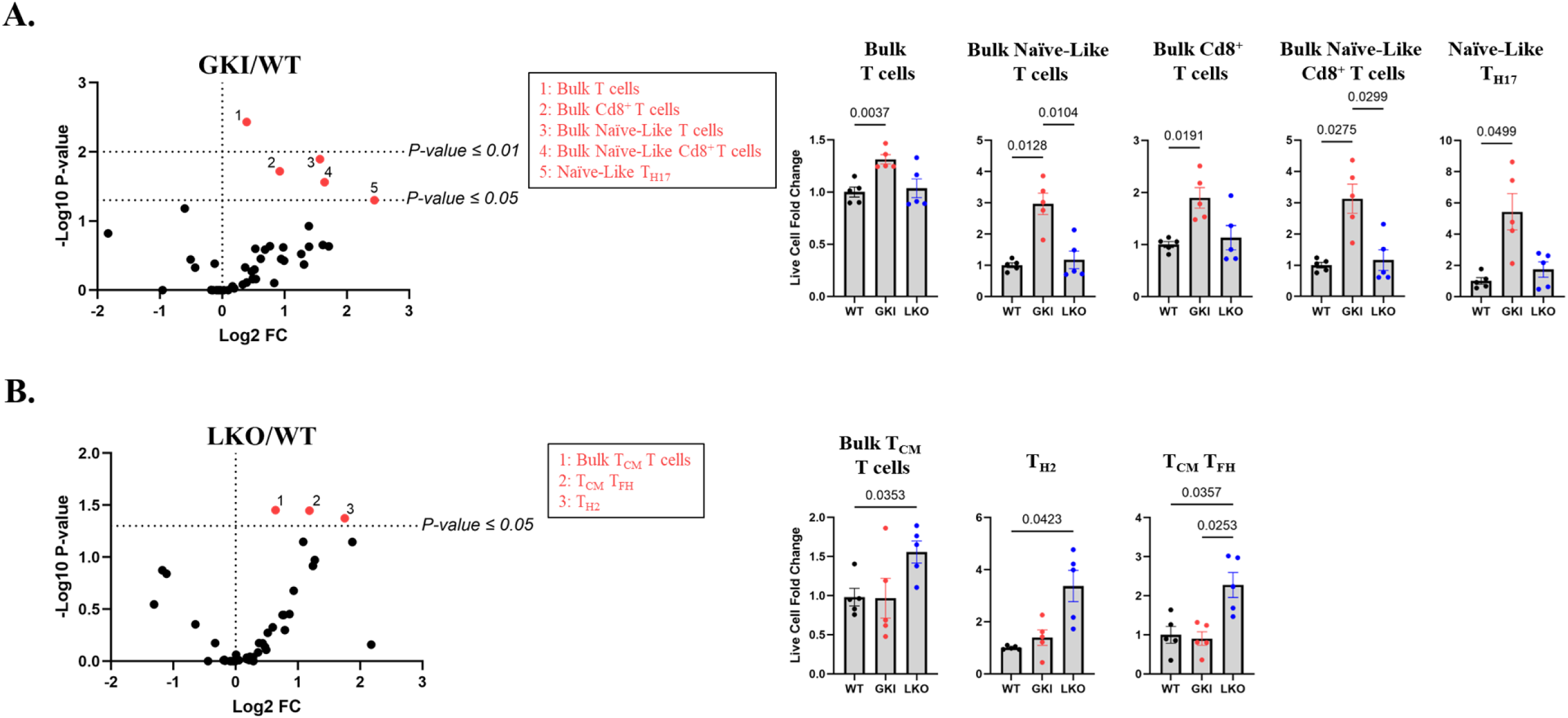
Bidirectional *Lrrk2* Disruptions Drive Divergent T cell Lineage Accumulations Within the Brain. Quantification via spectral flow cytometry of central T cell subset frequencies within the total live brain cells under basal conditions (N = 5 per genotype). Volcano plots map specific subset alterations in (**A.**) GKI/WT and (**B.**) LKO/WT configurations by charting the relationship between Log2 Fold Change and statistical significance (−Log10 *P-value*). Significantly altered subsets (*P-values ≤* 0.05 or *P-values ≤ 0.01*) are labeled in red and tracked to adjacent bar charts. Bar graphs plot individual biological replicates tracking Live Cell Fold Change (mean ± SEM). Exact *P-values* are annotated above brackets. For parametric data, statistical analysis was done using a Brown-Forsythe and Welch ANOVA with a Dunnett’s T3 multiple comparisons test. For nonparametric data, a Kruskal-Wallis test with a Dunn’s multiple comparison test was utilized.

As tissue-resident naïve-like cells are exceptionally rare, we applied a verification panel including Cxcr3 to distinguish true naïve cells from stem cell memory T cells (T_SCM_), which express Cd45ra^+^Cd62l^+^Cxcr3^+^ and are preferentially home in brain tissues (**Supplementary Table 4**; **Supplementary Figure 10** for gating strategy) [36–38]. As expected, most of the naïve-like T cell pool (62% of bulk T cells; 70.6% Cd8^+^ T cells) were identified as true T_SCM_ cells based upon a strong Cxcr3 expression, where both T_SCM_ bulk T cells and T_SCM_ Cd8^+^ T cells were significantly enriched in GKI brains (**Figure 5**). LKO brains did not have any significant changes in T_SCM_ T cells (**Figure 5**).

**Figure 5:**
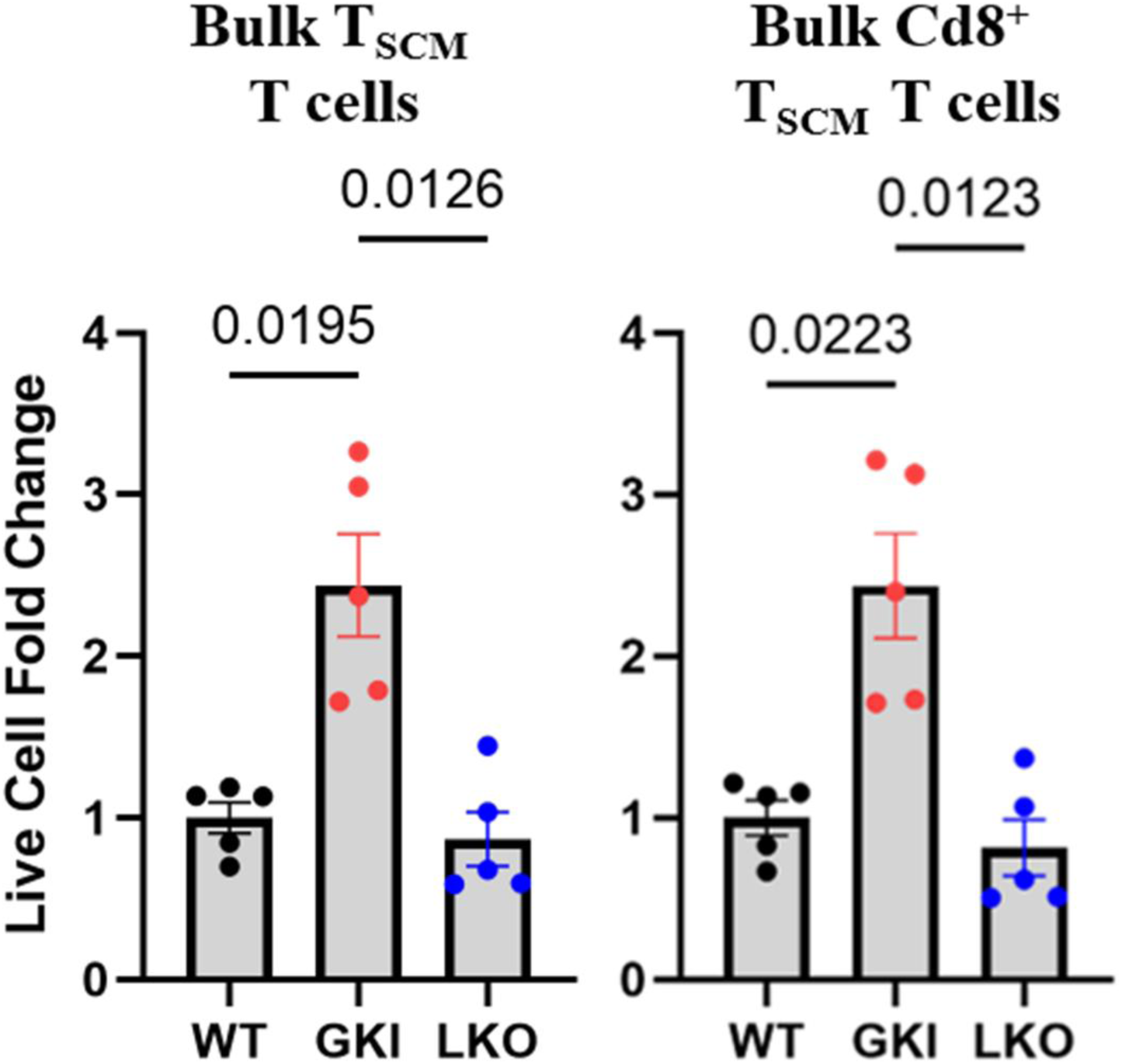
GKI Mice Exhibit Increased Expansion of Self-Renewing Stem Cell Memory T cells (T_SCM_) Within the Brain. Targeted immunophenotypic quantification of central tissue stem cell memory T cells (T_SCM_) classification of bulk T cells and bulk Cd8^+^ T cells were done with spectral flow cytometry (N = 5 per genotype). Bar graphs plot individual animal data points representing Live Cell Fold Change relative to WT controls (mean ± SEM). Exact *P-values* are annotated above brackets. Statistical analysis was done using a Brown-Forsythe and Welch ANOVA with a Dunnett’s T3 multiple comparisons test.

These results suggest that the GKI model promotes an influx or retention of self-renewing memory Cd8^+^ T cell populations and an overall higher T_H17_ cell population within the brain, whereas the LKO model appears to display a distinct type-2 like immune profile based upon high T_H2_ and T_FH_ cell populations alongside potential elevated immune surveillance[39, 40].

### Expansion of Peripheral Helper T cell Lineages is Driven by *Lrrk2* Genotypes

Utilizing the same flow cytometry immunophenotyping approach (**Supplementary Figure 4**; **Supplementary Table 4** for panel), we identified multiple changes in T cell subset populations within the periphery of our *Lrrk2* mouse models (see **Supplemental Figure 8B** for analysis of all immune subsets). In the periphery of GKI mice there is a significant expansion of conventional Cd4^+^ T cells (T_CON_), and more specifically an increase in those with effector memory (Cd45ra^−^ Cd62l^−^; T_EM_) and terminal effector (Cd45ra^+^Cd6l^−^; T_EF_) status (**Figure 6A**; **Supplementary Figure 11** for flow gating of significant findings). Further dissecting these subsets showed significant accumulations of both T_EM_ and T_EF_ variations of T_H1_, T_H2_, and T_H17_ (**Figure 6A**). LKO mice also showed a general increase in peripheral T_EM_ and T_EF_ Cd4^+^ T cells (**Figure 6B**). However, unlike the broad alterations in GKI mice, only the T_EM_ T_H17_ subset was significantly increased, while T_H1_ and T_H2_ subsets were not (**Figure 6B**). This indicates that alterations in *Lrrk2* genotype supports a systemic baseline bias toward highly active effector/memory Cd4^+^ T cell subsets, and more so in GKI than LKO mice, suggesting a fine programmatic balance of Lrrk2 activation is necessary to manage peripheral T cell homeostasis.

**Figure 6:**
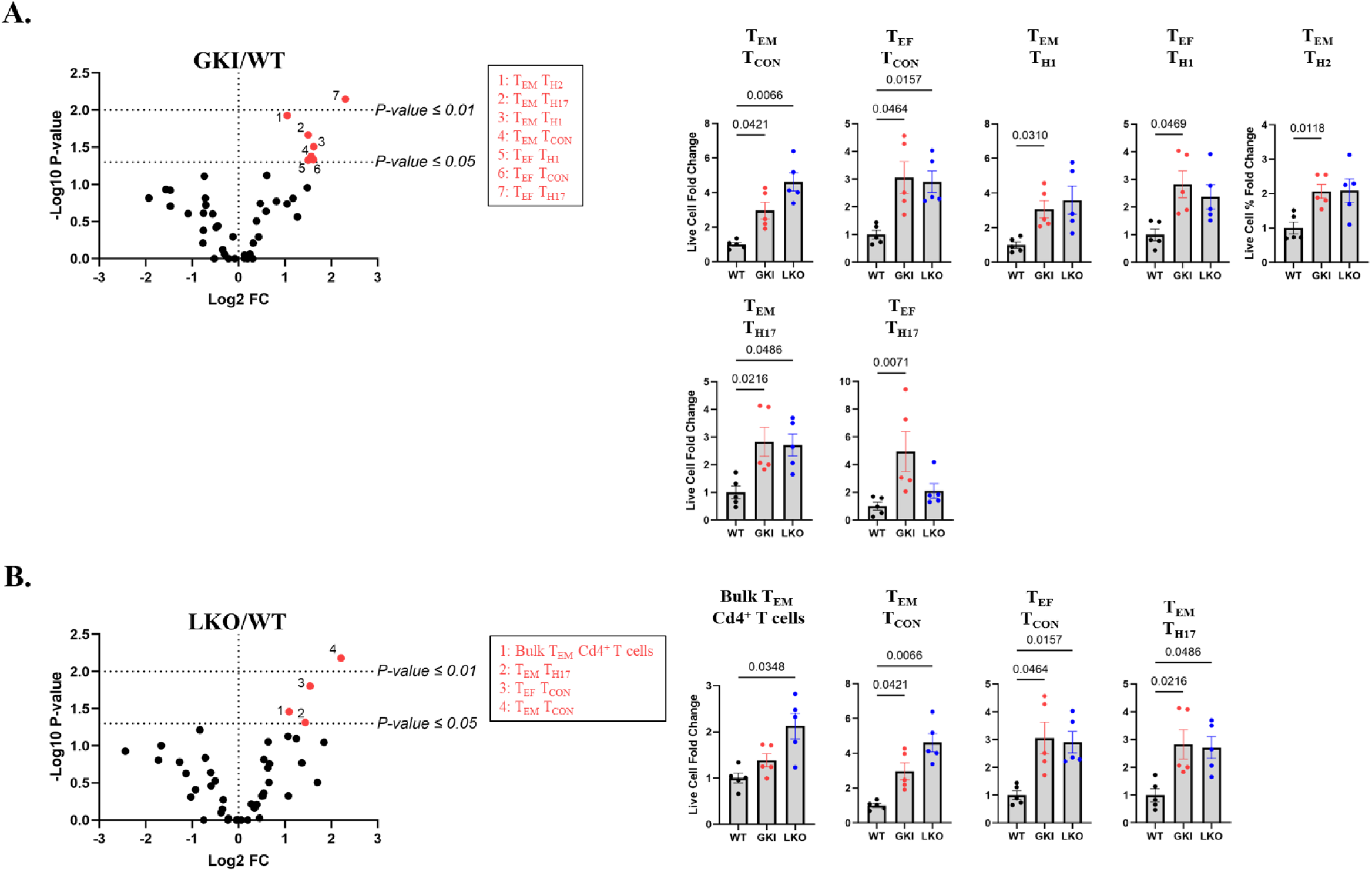
*Lrrk2* Genotype Influences a Systemic Baseline Expansion of Peripheral Helper T cell Lineages. Frequency alterations of peripheral conventional T cells (T_CON_) and effector configurations under basal conditions were determined through spectral flow cytometry (N =5 per genotype). Volcano plots map specific subset alterations in (**A.**) GKI/WT and (**B.**) LKO/WT configurations by charting the relationship between Log2 Fold Change and statistical significance (−Log10 *P-value*). Significantly altered subsets (*P-values ≤* 0.05 or *P-values ≤ 0.01*) are labeled in red and tracked to adjacent bar charts. Bar graphs plot individual biological replicates tracking Live Cell Fold Change (mean ± SEM). Exact *P-values* are annotated above brackets. For parametric data, statistical analysis was done using a Brown-Forsythe and Welch ANOVA with a Dunnett’s T3 multiple comparisons test. For nonparametric data, a Kruskal-Wallis test with a Dunn’s multiple comparison test was utilized.

### Acute *Ex Vivo* LPS Immune Challenge Exposes Genotype-Specific Transcript Suppression and Disrupts Inflammatory Priming

Following our immune cell baseline assessments within our mouse models, we evaluated the functional capacity of these isolated immune cell populations to an acute (6-hour) *ex vivo* immune challenge with LPS, dividing the samples evenly for transcriptomic (qPCR) and protein (flow cytometry) analyses. In an unbiased approach, similar to our previous qPCR experiments, we examined a variety of effector molecules in the total immune cell population: *Il1b* (Interleukin-1β), *Il2* (Interleukin-2), *Il4* (Interleukin-4), *Il6* (Interleukin-6), *Il10* (Interleukin-10), *Il13* (Interleukin-13), *Il17a* (Interleukin-17α), *Tnf* (Tumor Necrosis Factor), *Ifng* (Interferon γ), *Prf1* (Perforin), *Gzmb* (Granzyme b), and *Csf2* (Colony-Stimulated Factor 2; Granulocyte-Macrophage Colony-Stimulating Factor [GM-CSF]) (**Supplementary Table 1**). After stimulation, GKI brain immune transcripts demonstrated significant decreases in *Il4* and *Il6* expression, while having an increase in *Csf2* expression (**Figure 7A**). Interestingly, LKO brain immune cells revealed almost the opposite profile, where there was a significant increase of *Gzmb* gene expression (**Figure 7A**).

**Figure 7:**
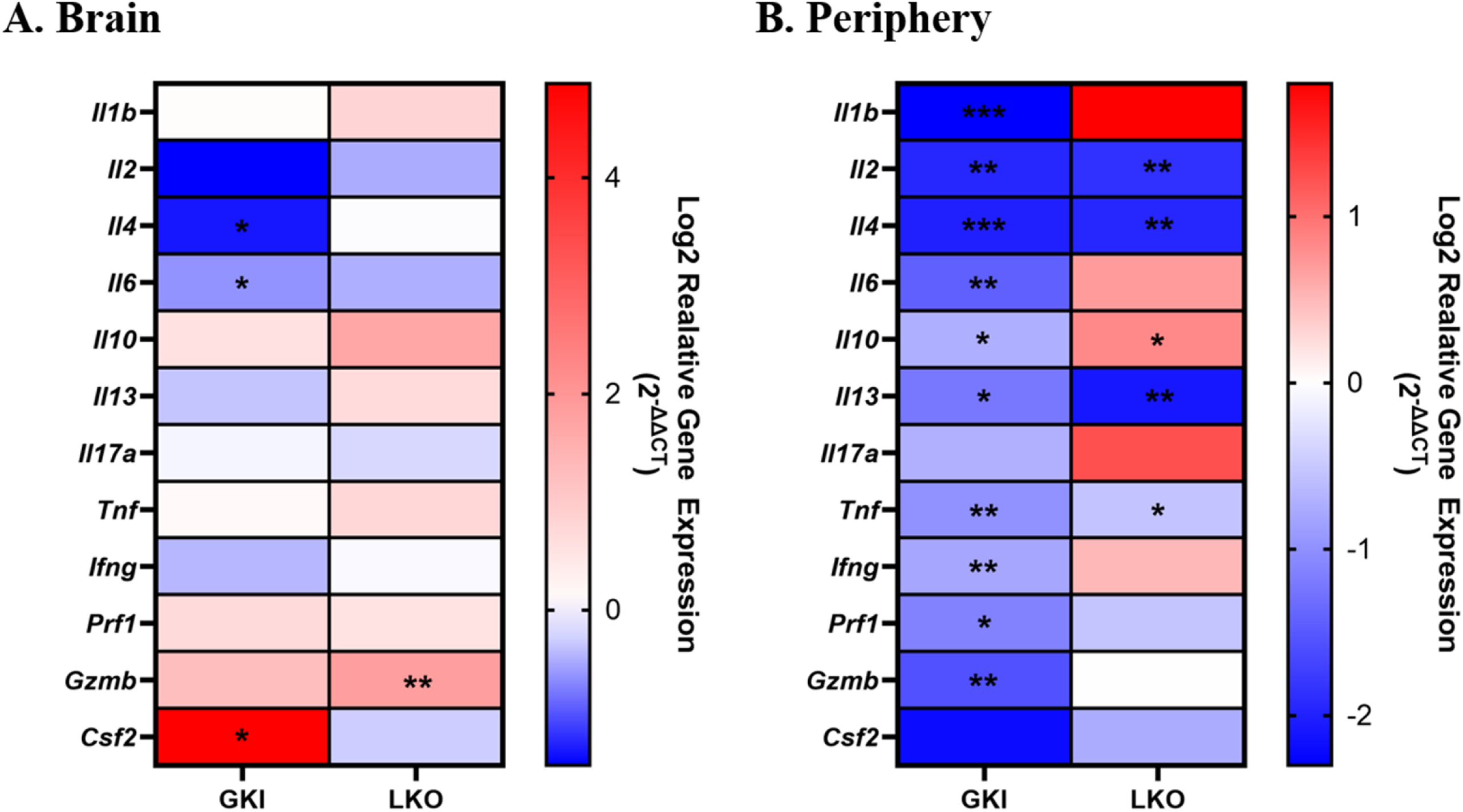
Acute *Ex Vivo* Lipopolysaccharide (LPS) Challenge Exposes Systemic Transcriptional Suppression and Impaired Inflammatory Priming in GKI Mice. (**A.**) Central (whole brain) and (**B.**) peripheral (PBMCs/splenocytes) gene expression profiles of effector molecule panel following a 6-hour acute *ex vivo* lipopolysaccharide (LPS) challenge (N = 5 per genotype). Data are presented as heatmaps of Log2 of relative gene expression (2^−ΔΔCT^) normalized to stimulated WT controls. Statistical analysis was done using a Brown-Forsythe and Welch ANOVA with a Dunnett’s T3 multiple comparisons test. A Benjamini-Hochberg FDR correction was applied for all *p-values* across all comparisons. Significance symbols: *: *P-value ≤ 0.05*; **: *P-value ≤ 0.01*; ***: *P-value ≤ 0.001*.

For the transcript levels of peripheral immune cells, it was an even greater surprise that the GKI mice overall demonstrated a severe, broad transcriptional suppression across almost all evaluated effector molecules (**Figure 7B**). This significant downregulation of gene expression included genes such as *Il1b*, *Il2*, *Il6*, *Il10, Il13*, *Tnf*, *Ifng*, *Prf1*, and *Gzmb* (**Figure 7B**). For the LKO peripheral immune cells, there was also significant downregulation of some of these genes (*Il2*, *Il4*, *Il13*, and *Tnf*), but not as pronounced as the GKI mice (**Figure 7B**). As within the brain, the LKO periphery also had a significant upregulation of *Il10* and *Gzmb* expression (**Figure 7B**). These gene dynamics reveal that the GKI genotype introduces systematic transcriptional dysregulation that severely limits rapid effector expression following immune activation, and impairment that is far less pronounced in the LKO model.

### Intracellular Effector Molecule Production is Paradoxically Stalled Across Both Mutant Lines After an Acute *Ex Vivo* LPS Immune Challenge

Next, we evaluated whether these significant alterations in transcription of the effector molecule genes would translate directly to intracellular protein production. Using our intracellular immunophenotyping panel in an unbiased manner (**Supplementary Table 3**; gating strategies in **Supplementary Figures 5 & 6**), we calculated the nMFI readings of all the intracellular effector molecules produced after an acute *ex vivo* LPS immune challenge of the mouse immune cells (**Figure 8**). Because resting immune cells produce trace amounts of intracellular cytokines that fall below the limits of flow cytometric detection, unstimulated baseline conditions yielded negligible intracellular staining across all genotypes (data not shown). Therefore, evaluating the true magnitude of effector molecule production necessitated an acute immune challenge.

**Figure 8:**
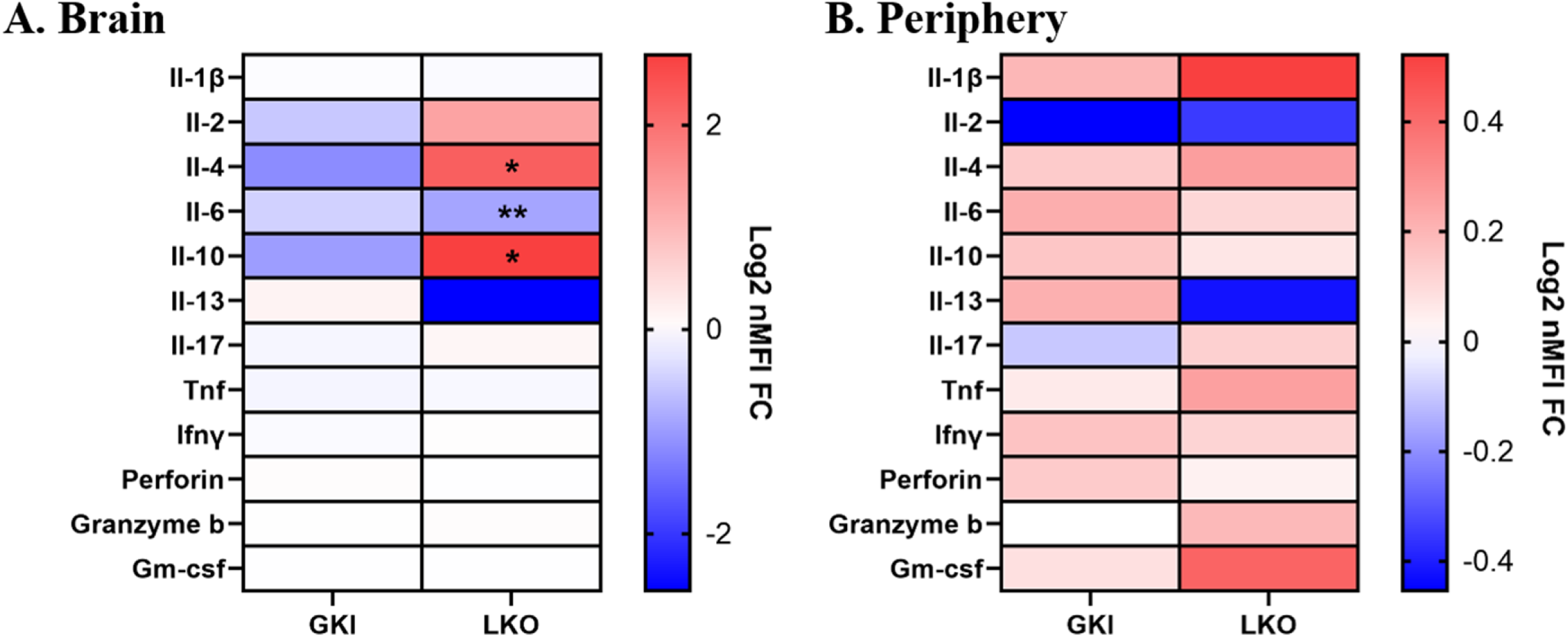
Acute *Ex Vivo* Lipopolysaccharide (LPS) Challenge Reveals a Post-Transcriptional Bottleneck Across Both *Lrrk2* Genotypes. (**A.**) Central (whole brain) and (**B.**) peripheral (PBMCs/splenocytes) relative protein expression profiles across an effector molecule spectral flow cytometry panel in overall live single cells following a 6-hour acute *ex vivo* lipopolysaccharide (LPS) challenge paired with a protein transport inhibitor (N = 5 per genotype). Heatmaps display the mean Log2 normalized median fluorescent intensity fold change (Log2 nMFI FC) relative to stimulated WT controls. For parametric data, statistical analysis was done using a Brown-Forsythe and Welch ANOVA with a Dunnett’s T3 multiple comparisons test. For nonparametric data, a Kruskal-Wallis test with a Dunn’s multiple comparison test was utilized. A Benjamini-Hochberg FDR correction was applied for all *p-values* across all comparisons. Significance symbols: *: *P-value ≤ 0.05*; **: *P-value ≤ 0.01*

In overall brain immune cells, GKI samples showed no significant protein production (**Figure 8A**). Interestingly, LKO brains showed significant protein increases for Il-4 and Il-10 alongside a significant decrease in Il-6 protein expression (**Figure 8A**). In the peripheral immune cells, while nMFI readings trended higher across several effector molecules in both GKI and LKO mice, there were no significant changes in overall protein production (**Figure 8B**).

This was further examined within single-cell subsets by evaluating iMFI values of intracellular molecule production across specific Cd4^+^ T cells, Cd8^+^ T cells, B cells, DCs, NK cells, and Cd11b^+^ phagocytes. Nevertheless, protein production in GKI mice compared to WT was largely unchanged (**Supplementary Figure 12**) as only peripheral NK cell perforin expression reached statistical significance (**Supplementary Figure 12B**). Conversely, LKO mice displayed altered intracellular molecule production, where stimulated central LKO Cd11b^+^ phagocytes (Tmem119^+^ only), Cd8^+^ T cells, B cells, and NK cells showed significantly reduced Il-6 and Tnf protein production (**Supplementary Figure 12A**). Conversely, Il-2 and Il-4 were significantly increased in central Cd11b^+^ phagocytes (Iba1^+^ and Tmem119^+^ populations), B cells, and DCs (**Supplementary Figure 12A**). Interestingly, peripheral LKO immune cells demonstrated a significant downregulation of Il-10 and Gm-csf within DCs, but an increase of Il-1b in Cd11b^+^ phagocytes (**Supplementary Figure 12B**). As the GKI model displayed flatline or only marginally elevated effector molecule production while their respective transcripts were heavily suppressed points to severe cellular dysregulation affecting protein/RNA synergy.

### Upregulation of the Inhibitory Surface Receptor Pd-1 Systematically on Multiple T cell Subsets Suggests a Chronic T cell Exhaustion Phenotype

To clarify the biological discrepancy previously observed earlier, where high baseline levels of membrane protein markers coincide with lower relative transcript levels, we aimed to understand the impact of stimulated effector molecule transcription being suppressed while effector protein production remains stalled. We investigated whether these models were exhibiting an exhaustion-like phenotype. Exhaustion phenotypes typically present as an accumulation of tissue or blood-resident cells that maintain low overall functionality due to a high baseline expression of inhibitory receptors such as Pd-1 and lymphocyte activation gene-3 (Lag-3)[41–46].

We first examined overall *Pdcd1* and *Lag3* expression levels in an unbiased manner in both central and peripheral immune systems (**Supplementary Figure 13**). Overall, there were no significant transcriptional changes across genotypes within the brain (**Supplementary Figure 13A**). However, in the periphery, *Pdcd1* expression was significantly reduced in GKI, while *Lag3* remained unchanged across all genotypes (**Supplementary Figure 13B**).

Because bulk transcriptional gene expression can mask cell-specific phenotypes, we utilized our flow cytometry immunophenotyping (gating strategies shown in **Supplementary Figure 14**) to determine protein expression of Pd-1 by iMFI calculation across various immune subsets (**Figure 9**; see **Supplemental Figure 15** for analysis of other immune subsets). In contrast with the suppression of gene expression, Pd-1 protein expression was significantly elevated on regulatory T cells (T_REG_) within GKI brains (**Figure 9A**). Peripheral protein expression of Pd-1 was also significant in the GKI mice on bulk Cd4^+^ T cells, specifically on T_H1_ and T_H2_ T cells (**Figure 9B**). Unexpectedly, in the periphery of the LKO and GKI mice, we observed similar significant increases of Pd-1 protein expression on the same Cd4^+^ T cells subsets (**Figure 9B**).

**Figure 9:**
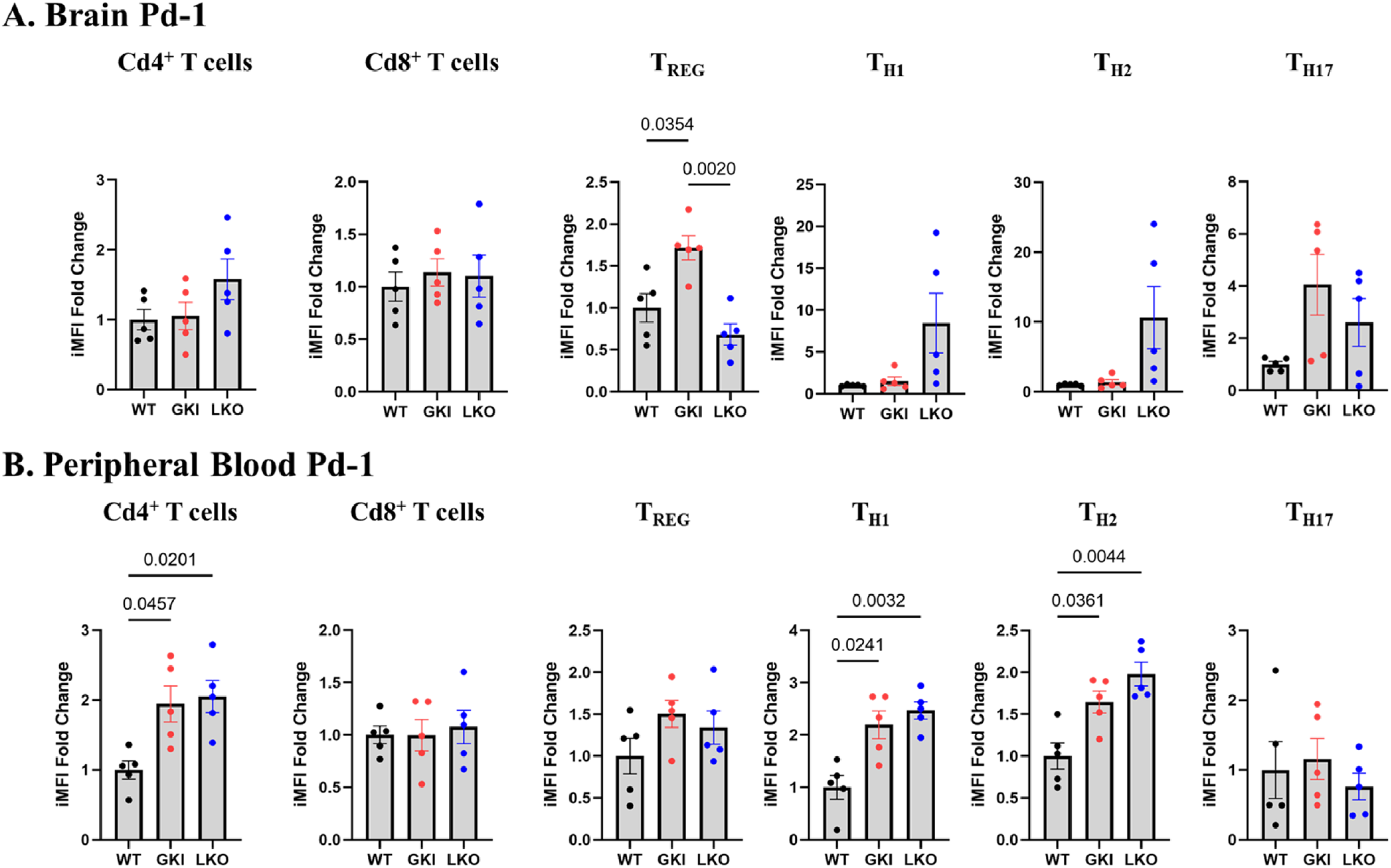
Surface Accumulation of the Inhibitory Receptor Pd-1 Corroborates a Phenotype of Chronic T cell Exhaustion. Integrated median fluorescent intensity (iMFI) calculations representing combined surface marker density and population frequency for Pd-1 expression across (**A.**) central (brain) and (**B.**) peripheral (PBMCs/splenocytes) T cell subsets under basal conditions using spectral flow cytometry (N = 5 per genotype). Bar graphs plot individual biological replicates tracking iMFI of Pd-1 (mean ± SEM). Exact *P-values* are annotated above brackets. For parametric data, statistical analysis was done using a Brown-Forsythe and Welch ANOVA with a Dunnett’s T3 multiple comparisons test. For nonparametric data, a Kruskal-Wallis test with a Dunn’s multiple comparison test was utilized.

This pronounced surface accumulation of Pd-1 on central and peripheral T cells and the flattened functional effector outputs with suppressed transcriptional profiles suggest an exhaustion-like state induced in the GKI condition and, to a lesser extent, in the LKO model. Overall, the balance of Lrrk2 activity is crucial in maintaining immune homeostasis.

## DISCUSSION

Pathogenic mutations in *LRRK2* (p.G2019S)[5–7], vacuolar protein sorting 35 (*VPS35* [p.D620N])[47–49], and Rab GTPase 32 (*RAB32* [p.S71R])[20, 50, 51] cause PD, and are all associated with increased LRRK2 kinase activity. All three proteins play important roles in innate immunity and pathogen defense, mediated via the MiT transcription factor family, and LRRK2 and RAB32 are especially enriched within myeloid cells[8, 10, 52–54]. To understand how LRRK2 influences immunoregulation in both central and peripheral immunity, we combined multi-parameter spectral flow cytometry immunophenotyping with quantitative transcriptomics to profile the immune consequences of *Lrrk2* overactivation (GKI mice) and genetic deletion (LKO mice). Our findings demonstrate *Lrrk2* is an essential mediator of immunological homeostasis, specifically in T cell biology, where bidirectional alteration in *Lrrk2* (gain-of-function or complete loss) produces distinct, but convergent, patterns of immune exhaustion.

*Lrrk2* appears to play a role in immune subset differentiation and homing across multiple immune cell types, at baseline, and particularly with T cell subsets. While most *Lrrk2* research has focused on peripheral blood, splenic studies, and central innate immunity (microglia) in humans and mice[8, 10, 52–54], few investigations have examined how Lrrk2 p.G2019S or complete *Lrrk2* knockout affects both central and peripheral immunity within the same animal. Within the brain, the GKI mice had increases in both bulk Cd4^+^ and Cd8^+^ T cells and specifically T_H17_ Cd4^+^ T cells, while LKO mice had distinct expansions of T_H2_ and T_FH_ Cd4^+^ subsets. These phenotypes indicate that the p.G2019S mutation broadly enhances overall T cell subset expansion, while loss of *Lrrk2* leads to increases in specific T cell subsets within central immunity. In the periphery, both models demonstrated increased effector and effector memory Cd4^+^ T cell populations, indicating a shared peripheral immune activation phenotype despite divergent central signatures. These observations confirm that at baseline levels of inflammation, Lrrk2 activity appears necessary to maintain homeostatic T cell distributions between both central and peripheral immunity.

The cellular plasticity and memory differentiation capacity of T cells are essential elements of adaptive immune function[55–57]. As such, we included Cd45ra^+/−^ and Cd62l^+/−^ immunophenotyping to recognize these diverse populations of memory and naïve T cells. Recent investigations into T cell memory have focused on neurodegeneration and brain injury[58–61]. Most studies suggest memory Cd8^+^ T cells are more prevalent in the brain, while memory Cd4^+^ T cells dominate the periphery, which we confirm in our *Lrrk2* mouse models at baseline. We also discovered a significant expansion of self-renewing, antigen-experienced early memory T cells (T_SCM_) in GKI mouse brain that was significantly less within LKO and WT controls. T_SCM_ T cells express a naïve-like (Cd45ra^+^Cd62l^+^) phenotype along with Cxcr3^+^ expression[62–64] to help guide memory T cells within brain tissue but have not been thoroughly examined in neurodegenerative studies. The co-occurrence of elevated T_SCM_ populations alongside expanded peripheral T_EM_ and T_EF_ T cell populations in the same GKI animals indicates a genetic predisposition to accelerated immune aging/exhaustion driven by constitutive Lrrk2 kinase activity[65, 66]. We postulate Lrrk2 controls memory differentiation, but further *ex vivo* and *in vivo* assays in naïve T cell populations are needed to validate how Lrrk2 kinase activation alters memory fate decisions.

One central finding is the pronounced dissociation between gene expression and protein translation, while predominantly observed in GKI periphery, is apparent in both mouse models. Discordant rates of mRNA transcription and surface protein expression are a feature of chronically activated and exhausted cells[67–69]. In GKI mice, peripheral immune transcripts were broadly suppressed while flow cytometry revealed expanded T cell populations with elevated surface marker expression, which closely aligns with an exhausted immune profile. This is also evident in the acute *ex vivo* LPS challenge, where GKI peripheral immune cells underwent near-global transcript suppression and stalled protein output of effector molecules. LKO immune cells also displayed this phenomenon, but to a lesser extent, demonstrating bidirectional disruption of *Lrrk2* can promote an immune exhausted state. This chronic state likely reflects the downstream consequences of impaired endolysosomal trafficking and macroautophagy caused by the dysregulation of Lrrk2 kinase activity and aberrant Rab GTPase phosphorylation[70–77].

Our exhaustion hypothesis was directly supported by Pd-1 surface expression profiling across immune subsets, which is a classical indicator of general immune cell exhaustion[44–46]. In the brains of the GKI mice, Pd-1 expression was significantly elevated on T_REG_ cells, implying an impairment of central T cell suppression. In the periphery, there were significant increases of Pd-1 found on bulk Cd4^+^ T cells, specifically in T_H1_ and T_H2_ subsets in both GKI and LKO mice. The convergence of elevated peripheral Pd-1 across both hyperactive and knockout models of *Lrrk2*, despite their divergent central phenotypes and transcriptional profiles, provide strong evidence for a loss-of-homeostasis model in which Lrrk2 activity must be tightly regulated to prevent exhaustion. Excess Lrrk2 kinase activity disrupts lysosomal function and mitophagy[72, 74, 75, 77], promoting protein accumulation and mitochondrial stress, while loss of Lrrk2 activity can cause disruption of basal mitophagy and autophagic flux[74, 78, 79]. While the GKI and LKO models appear to be in an exhaustion-like state, our immunophenotyping reveals they take divergent pathways to get there. Our studies demonstrate that GKI predominantly accumulate T_H1_, T_H17_, and Cd8^+^ T cell subsets, reflecting a highly proinflammatory and cytotoxic profile. Chronic hyperactivation of these specific inflammatory pathways naturally triggers exhaustion mechanisms as a compensatory fail-safe to prevent severe immunopathology[80, 81]. Conversely, it appears LKO mice exhibit expansions in T_H2_ and T_FH_ subsets. While traditionally associated with humoral immunity, dysregulation of the expansions of these cells paired with a loss of Lrrk2 activation controlling autophagic and metabolic regulation can similarly drive T cells into an exhaustion-like state, thus stalling functional output[9, 41, 74]. Therefore, the functional convergence we observed demonstrates that while Lrrk2 overactivity drives exhaustion through chronic inflammatory burnout, complete loss of Lrrk2 drives a similar unresponsiveness likely through metabolic dysregulation and subset skewing, but further investigation is required.

Several limitations of this study should be acknowledged. Immune cell populations were assessed without an *in vivo* antigenic challenge, reflecting a deliberate choice to characterize the purely genetic effects of *Lrrk2* status at baseline. Future studies using *in vivo* LPS or α-syn fibril injection models may determine how these baseline alterations manifest under physiological inflammatory conditions. The LKO models do not fully replicate the pharmacological effects of MLi-2 kinase inhibition, as we originally hypothesized that the LKO model would be able to replicate previous studies that utilized MLi-2[47, 82, 83]. Nevertheless, as demonstrated, complete loss of Lrrk2 still causes subtle immune cell dysregulation. Future work with cell type-specific MLi-2 application would help elucidate how kinase suppression produces an immune phenotype distinct from a complete knockout. While *ex vivo* LPS was used to examine intracellular effector molecule production, concentration and incubation time were minimized to remain as close to baseline as possible. Thus, future studies might examine dose and time-dependent immune challenge responses. Further investigation of lysosomal/endosomal processes and autophagic flux, focused on exhaustion markers in isolated naïve, active, memory, and exhausted T cell subsets, are also needed. To understand the mechanistic relationship between Lrrk2 kinase activity, T cell exhaustion, microglial activation and the insidious loss of nigral neurons in PD, is a *sine qua non* to develop therapeutic interventions.

## CONCLUSIONS

In summary, this study establishes that Lrrk2 kinase activity is an essential regulator of T cell homeostasis in both the central and peripheral immune systems and shows bidirectional dysregulation of this activity drives T cells towards functional exhaustion. Lrrk2 p.G2019S and *Lrrk2* knockout each disrupt the balance between T cell activation, memory differentiation, and effector function through mechanistically distinct pathways. These findings have direct implications for clinical development, as therapeutic strategies must carefully titrate LRRK2 inhibition to avoid driving the immune system from one dysfunctional extreme to another. Immune cell function should be carefully monitored when targeting LRRK2 kinase activity in patients with PD.

## LIST OF ABBREVIATIONS

PD: Parkinson’s disease
LRRK2/Lrrk2: Leucine rich-repeat 2
WT: Wild-type C57BL/6J
GKI: C57BL/6J-B6.Cg-*Lrrk2^tm1.1Hlme^*/J
LKO: C57BL/6-*Lrrk2^tm1Mjfa^*/J
qPCR: Quantitative polymerase chain reaction
IACUC: Institutional Animal Care and Use Committee
NIH: National Institute of Health
HBSS: Hank’s Balanced Salt Solution
FBS: Fetal bovine serum
PBMCs: Peripheral blood mononuclear cells
cDNA: Complementary DNA
FMO: Fluorescent minus one
cDMEM/F-12: Complete Dulbecco’s Modified Eagle Medium/Nutrient Mixture F-12
LPS: Lipopolysaccharide
FCS: Flow cytometry standard
MFI: Median fluorescent intensity
nMFI: Normalized MFI
iMFI: Integrated MFI
MIFlowCyt: Minimum Information about a Flow Cytometry Experiment
tSNE: T-distributed stochastic neighbor embedding
Cd3e: Cluster of differentiation 3 epsilon
Cd8a: Cluster of differentiation 8 alpha
Cd4: Cluster of differentiation 4
Itgax/Cd11c: Integrin alpha x/Cluster of differentiation 11c
DCs: Dendritic cells
Ncr1/Nkp46: Natural cytotoxicity triggering receptor 1/Nkp46
NK cells: Natural Killer Cells
Cd22: Cluster of differentiation 22
Itgam/Cd11b: Integrin alpha m/Cluster of differentiation 11b
Tmem119: Transmembrane protein 119
Aif1/Iba1: Allograft inflammatory factor 1/Ionized calcium binding adapter molecule 1
Gfap: Glial fibrillary acidic protein
S100b: S100 calcium-binding protein b
Cd183/Cxcr3: Cluster of differentiation 183/Chemokine receptor Cxcr3
T_H1_: T helper 1
Cd45ra: Cluster of differentiation 45ra/Protein tyrosine phosphatase, receptor type, C
Cd62l: Cluster of differentiation 62l/L-selectin
Cd196/Ccr6: Cluster of differentiation 196/Chemokine receptor 6
T_H17_: T helper 17
Pd-1/Pcd1/Cd279: Programmed cell death protein 1/Cluster of differentiation 279
T_FH_: T follicular helper
Cd185/Cxcr5: Cluster of differentiation 185/C-X-C chemokine receptor type 5
Cd25/Il-2ra: Cluster of differentiation 25/Interleukin-2 receptor alpha
Cd127/Il-17ra: Cluster of differentiation 127/Interluekin-17 receptor alpha
Cd86/B7-2: Cluster of differentiation 86/B7-2
T_CM_: Central memory T cell
T_H2_: T helper 2
T_SCM_: Stem cell memory T cell
T_CON_: Cd4^+^ conventional T cell
T_EM_: Effector memory T cell
T_EF_: Effector T cell
Il1b: Interleukin-1 beta
Il2: Interleukin-2
Il4: Interleukin-4
Il6: Interleukin-6
Il10: Interleukin-10
Il13: Interleukin-13
Il17a: Interluekin-17 alpha
Tnf: Tumor necrosis factor
Ifng: Interferon gamma
Prf1: Perforin
Gzmb: Granzyme b
Csf2/Gm-csf: Colony-stimulated factor 2/Granulocyte-macrophage colony-stimulating factor
Lag-3: Lymphocyte activation gene-3
T_REG_: Regulatory T cell
VPS35: Vacuolar protein sorting-associated protein 35
RAB322: Rab GTPase 32

## DECLARATIONS

### Ethics Approval and Consent to Participate

Not Applicable

### Consent for Publication

Not Applicable

### Availability of Data and Materials

The datasets generated and/or analyzed during the current study are available in the BioRxiv repository as a pre-print (doi:XXX). The datasets used and/or analyzed are also available from the corresponding author on reasonable request. All data generated or analyzed during this study are included in this published article and its supplementary information files.

### Authors’ Contributions

Conceptualization: RCS, MJF

Methodology: RCS, SCW, JF, IBD

Investigation: RCS

Formal analysis: RCS

Visualization: RCS

Funding acquisition: MJF

Project administration: MJF

Resources: MJF

Supervision: MJF

Writing – original draft: RCS, JF, IBD, MJF

Writing – review & editing: RCS, SCW, JF, IBD, MJF

## Acknowledgments

The Farrer lab would like to thank Drs. Melissa A. Maczis and Dylan T. Guenther for their technical support. We appreciate Dr. Malu Tansey and her lab for their help in experimental design.

