## Supplementary Figures and Tables for "Bidirectional disruption of Lrrk2 function drives T cell dysregulation and an exhaustion-like immune response"

**SUPPLEMENTARY FIGURES AND TABLES FOR SHARP ET AL., BIDIRECTIONAL DISRUPTION OF LRRK2 FUNCTION DRIVES T CELL DYSREGULATION AND AN EXHAUSTION-LIKE IMMUNE RESPONSE**

### SUPPLEMENTARY FIGURES AND LEGENDS

**Supplementary Figure 1: Representation of Flow Cytometry Gating Strategy for Median Fluorescent Intensity (MFI) Tracking of Immune Subset Markers in the Brain Single-Cell Suspensions**

Flow cytometry immunophenotyping of single-cell suspensions isolated from brains of WT, GKI, and LKO mouse models (N = 5 per genotype). All gate placements and boundary configurations were established using unstained controls, single color controls, and fluorescence-minus-one (FMO) controls. A comprehensive inventory of antibodies and fluorophores is provided in **Supplementary Table 2.** To visualize the gating workflow, raw FCS files from all biological replicates across all cohorts were concatenated into a single master file (15 files total), (**A.**). Sequential manual gating strategy used to isolate single, live mononuclear cells. (**B.**). Deconvolution of the master dataset back into individual cohorts (WT, GKI, LKO) via SampleID keyword tracking, followed by targeted gating to evaluate expression levels for the following surface markers: Cd3$^{+}$, Cd8a$^{+}$, Cd4$^{+}$, Cd127$^{+}$ (Il-7ra), Cd25$^{+}$ (Il-2ra), Pd-1$^{+}$, Cd185$^{+}$ (Cxcr5), Cd183$^{+}$ (Cxcr3), Cd194$^{+}$ (Ccr4), Cd196$^{+}$ (Ccr6), Cd45ra$^{+}$, Cd62l$^{+}$ (L-selectin), Cd11c$^{+}$ (Itgax), Nkp46$^{+}$ (Ncr1), Cd22$^{+}$, Cd11b$^{+}$ (Itgam), Tmem119$^{+}$, Iba1$^{+}$ (Aif1), Cd14$^{+}$, Cd68$^{+}$, Cd86$^{+}$ (B7-2), Cd163$^{+}$, Gfap$^{+}$, and S100b$^{+}$.

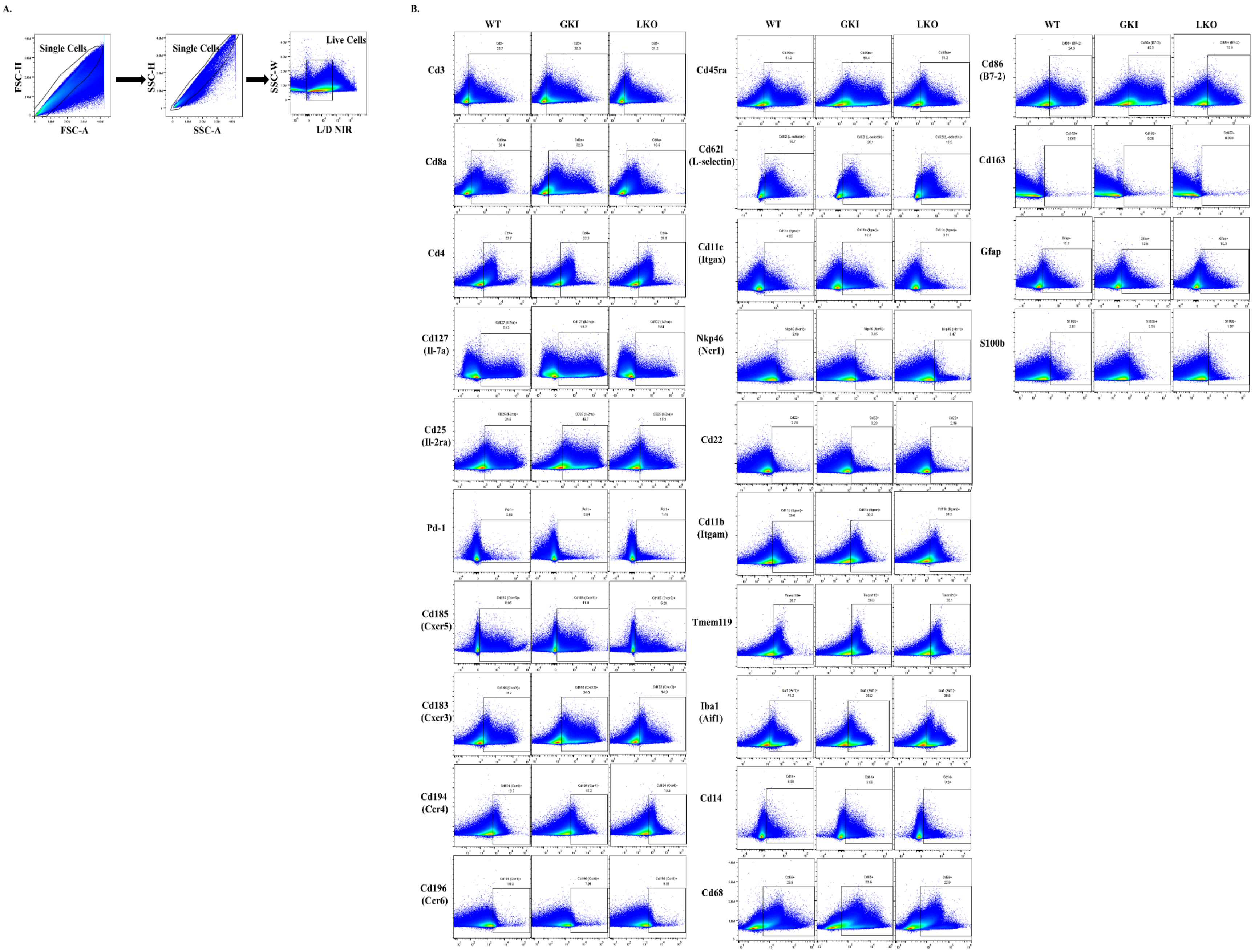
A.
Single Cells
FSC-H
FSC-A
Single Cells
SSC-H
SSC-A
Live Cells
SSC-W
L/D NIR
B.
WT
GKI
LKO
Cd3
Cd8a
Cd4
Cd127 (Il-7a)
Cd25 (Il-2ra)
Pd-1
Cd185 (Cxcr5)
Cd183 (Cxcr3)
Cd194 (Ccr4)
Cd196 (Ccr6)
Cd45ra
Cd62l (L-selectin)
Cd11c (Itgax)
Nkp46 (Ncr1)
Cd22
Cd11b (Itgam)
Tmem119
Iba1 (Aif1)
Cd14
Cd68
Cd86 (B7-2)
Cd163
Gfap
S100b

**Supplementary Figure 2: Representation of Flow Cytometry Gating Strategy for Median Fluorescent Intensity (MFI) Tracking of Immune Subset Markers in Peripheral blood (PBMCs) and Splenocyte Single-Cell Suspensions**

Flow cytometry immunophenotyping of single-cell suspensions isolated from combined peripheral blood mononuclear cells (PBMCs) and splenocytes of WT, GKI, and LKO mouse models (N = 5 per genotype). All gate placements and boundary configurations were established using unstained controls, single color controls, and fluorescence-minus-one (FMO) controls. A comprehensive inventory of antibodies and fluorophores is provided in **Supplementary Table 2.** To visualize the gating workflow, raw FCS files from all biological replicates across all cohorts were concatenated into a single master file (15 files total), (**A.**). Sequential manual gating strategy used to isolate single, live mononuclear cells. (**B.**). Deconvolution of the master dataset back into individual cohorts (WT, GKI, LKO) via SampleID keyword tracking, followed by targeted gating to evaluate expression levels for the following surface markers: Cd3$^+$, Cd8a$^+$, Cd4$^+$, Cd127$^+$ (Il-7ra), Cd25$^+$ (Il-2ra), Pd-1$^+$, Cd185$^+$ (Cxcr5), Cd183$^+$ (Cxcr3), Cd194$^+$ (Ccr4), Cd196$^+$ (Ccr6), Cd45ra$^+$, Cd62l$^+$ (L-selectin), Cd11c$^+$ (Itgax), Nkp46$^+$ (Ncr1), Cd22$^+$, Cd11b$^+$ (Itgam), Tmem119$^+$, Iba1$^+$ (Aif1), Cd14$^+$, Cd68$^+$, Cd86$^+$ (B7-2), Cd163$^+$, Gfap$^+$, and S100b$^+$.

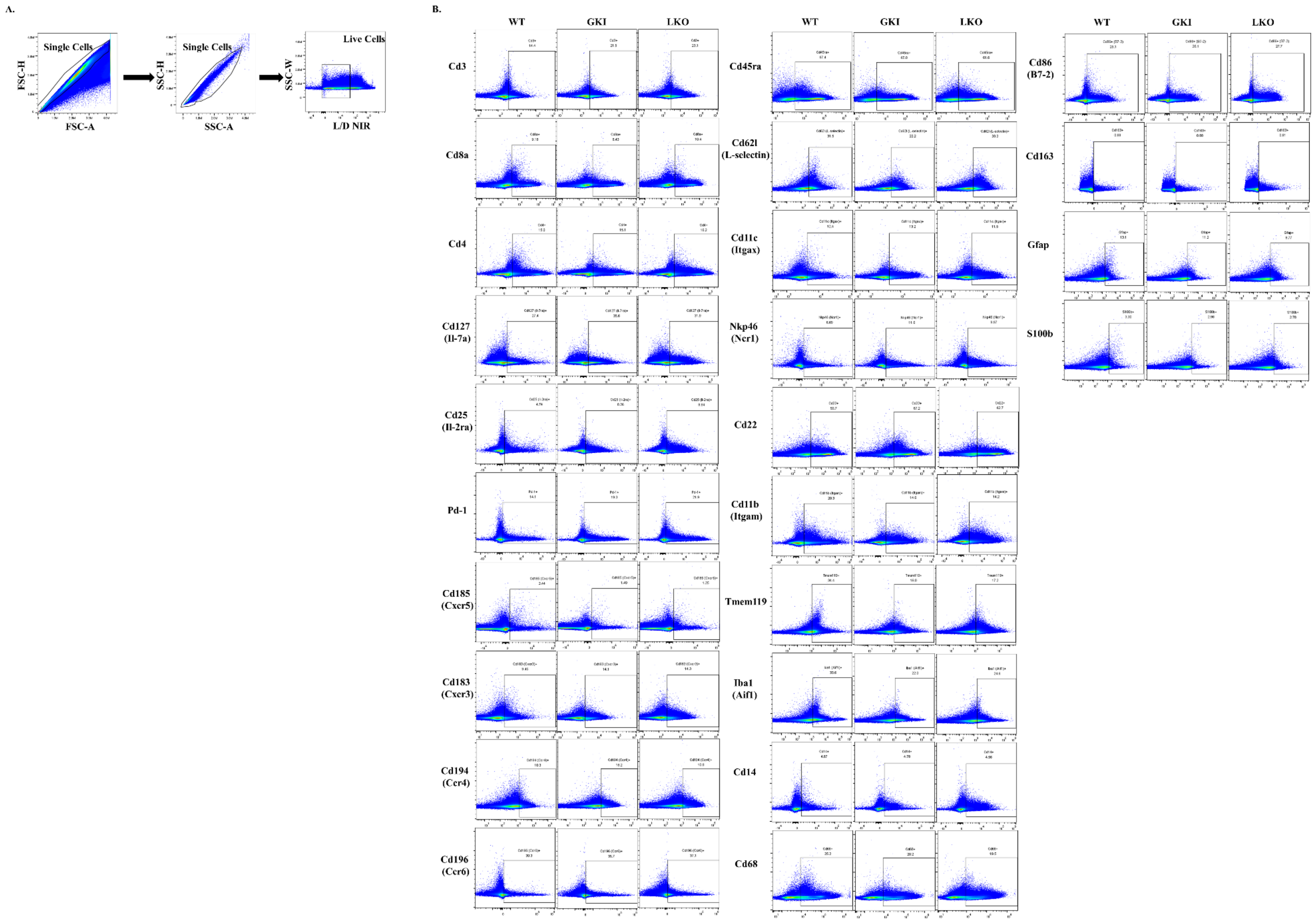
A.
Single Cells
FSC-H
FSC-A
Single Cells
SSC-H
SSC-A
Live Cells
SSC-W
L/D NIR
B.
WT
GKI
LKO
Cd3
Cd8a
Cd4
Cd127 (Il-7a)
Cd25 (Il-2ra)
Pd-1
Cd185 (Cxcr5)
Cd183 (Cxcr3)
Cd194 (Ccr4)
Cd196 (Ccr6)
Cd45ra
Cd62l (L-selectin)
Cd11c (Itgax)
Nkp46 (Ncr1)
Cd22
Cd11b (Itgam)
Tmem119
Iba1 (Aif1)
Cd14
Cd68
Cd86 (B7-2)
Cd163
Gfap
S100b

**Supplementary Figure 3: Flow Cytometry Manual Gating Strategy for Distinct Immune Cell Subset Population Identification Within Brain Tissue**

Representative gating tree for structural population profiling of central (brain) immune system (N = 5 per genotype). Initial gates were calibrated via unstained, single-color, and fluorescence-minus-one (FMO) controls, utilizing antibody combinations specifically in **Supplementary Table 2**. All 15 individual sample FCS files were concatenated to construct a unified template for upstream gating of live single cells. Downstream gates were sequentially applied to resolve complex multi-marker immune phenotypes. Lineage definitions, gating hierarchies, and peer-reviewed classification references for each identified subset are detailed in **Supplementary Table 4**.

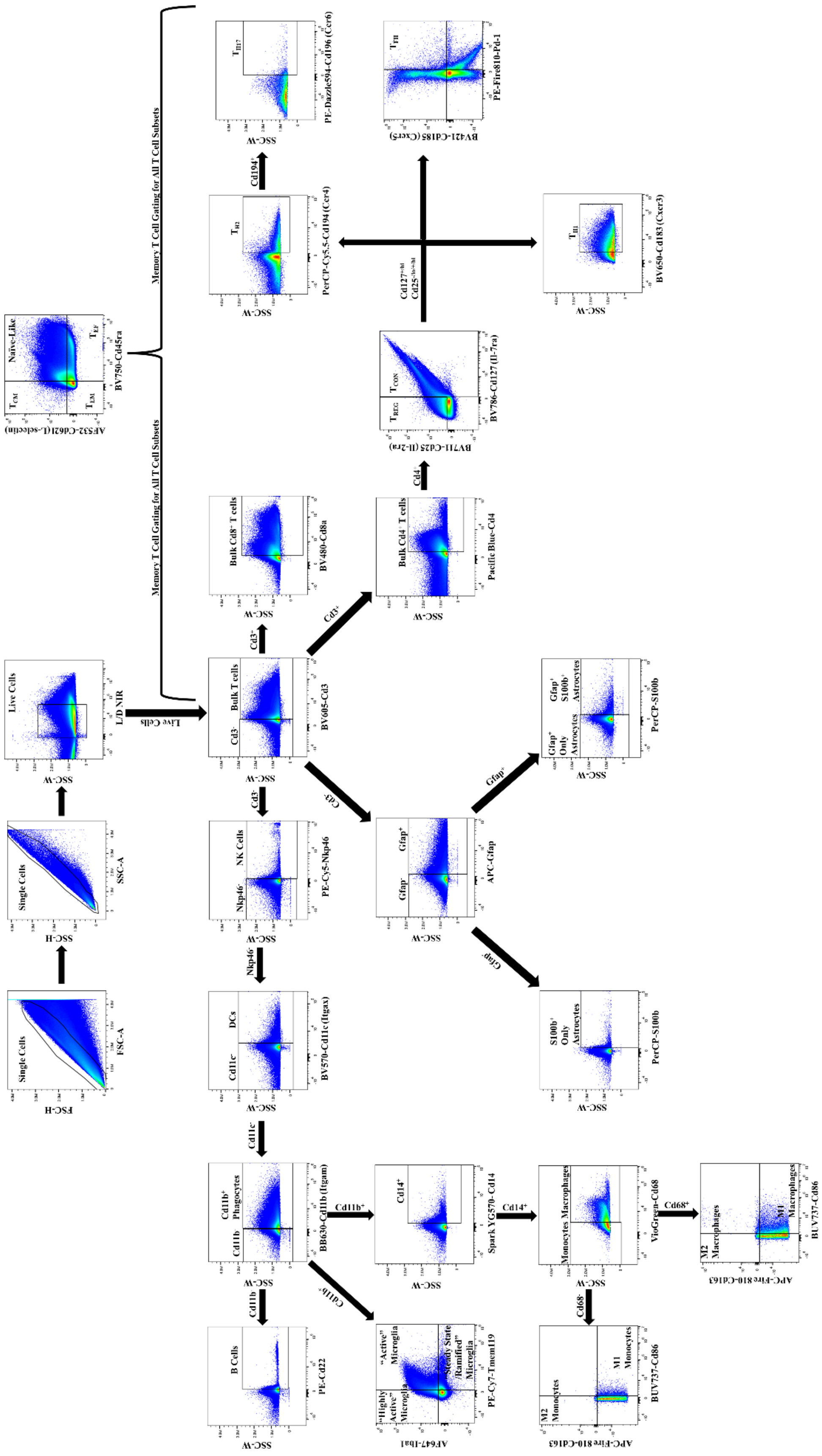

Single Cells
FSC-H
FSC-A
Single Cells
SSC-H
SSC-A
Live Cells
SSC-W
L/D NIR
Live Cells
Memory T Cell Gating for All T Cell Subsets
Memory T Cell Gating for All T Cell Subsets
Naïve-Like
TCM
TEM
TEF
AF532-Cd62l (L-selectin)
BV750-Cd45ra
Cd3-
Bulk T cells
BV605-Cd3
Cd3+
Bulk Cd8+ T cells
BV480-Cd8a
TH2
PerCP-Cy5.5-Cd194 (Ccr4)
Cd194+
TH17
PE-Dazzle594-Cd196 (Ccr6)
Cd3+
Bulk Cd4+ T cells
Pacific Blue-Cd4
Cd4+
TREG
TCON
BV711-Cd25 (Il-2ra)
BV786-Cd127 (Il-7ra)
TFH
BV421-Cd185 (Cxcr5)
PE-Fire810-Pd-1
TH1
BV650-Cd183 (Cxcr3)
Cd3-
Nkp46-
NK Cells
PE-Cy5-Nkp46
Nkp46-
Cd11c-
DCs
BV570-Cd11c (Itgax)
Cd3-
Gfap-
Gfap+
APC-Gfap
Gfap+
Gfap+ Only Astrocytes
Gfap+ S100b+ Astrocytes
PerCP-S100b
Gfap-
S100b+ Only Astrocytes
PerCP-S100b
Cd11c-
Cd11b
Cd11b+ Phagocytes
BB630-Cd11b (Itgam)
Cd11b+
Cd14+
Spark YG570-Cd14
Cd14+
Monocytes
Macrophages
VioGreen-Cd68
Cd68+
M2 Macrophages
M1 Macrophages
APC-Fire 810-Cd163
BUV737-Cd86
Cd11b
B Cells
PE-Cd22
Cd11b-
"Highly Active" Microglia
"Active" Microglia
"Steady State /Ramified" Microglia
AF647-Iba1
PE-Cy7-Tmem119
Cd68-
M2 Monocytes
M1 Monocytes
APC-Fire 810-Cd163
BUV737-Cd86

**Supplementary Figure 4: Flow Cytometry Manual Gating Strategy for Distinct Immune Cell Subset Population Identification in Peripheral blood (PBMCs) and Splenocyte Single-Cell Suspensions**

Representative gating tree for structural population profiling of peripheral (PBMCs/splenocytes) immune system (N = 5 per genotype). Initial gates were calibrated via unstained, single-color, and fluorescence-minus-one (FMO) controls, utilizing antibody combinations specifically in **Supplementary Table 2**. All 15 individual sample FCS files were concatenated to construct a unified template for upstream gating of live single cells. Downstream gates were sequentially applied to resolve complex multi-marker immune phenotypes. Lineage definitions, gating hierarchies, and peer-reviewed classification references for each identified subset are detailed in **Supplementary Table 4**.

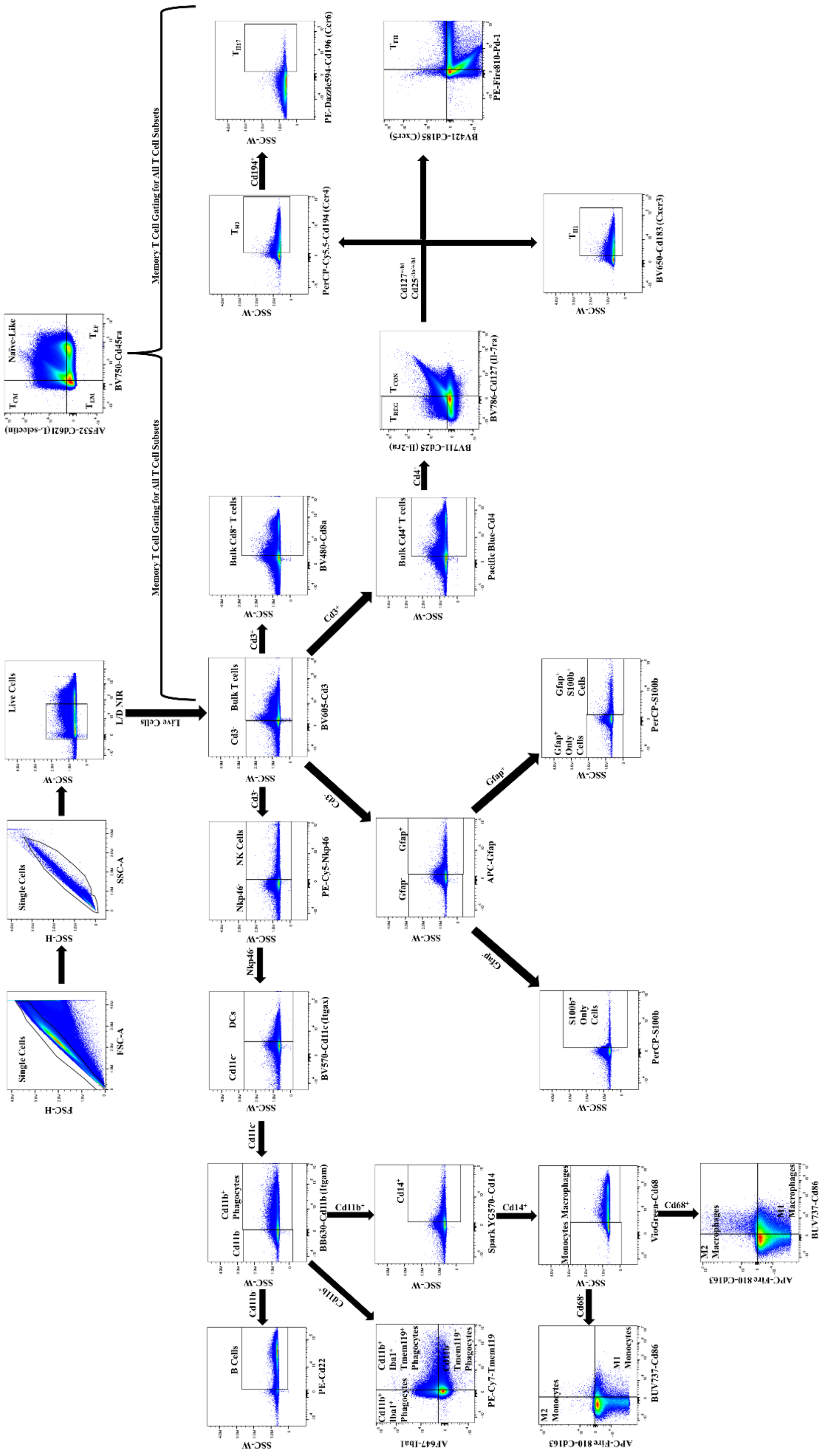

Single Cells
FSC-A
FSC-H
Single Cells
SSC-A
SSC-H
Live Cells
L/D NIR
SSC-W
Live Cells
Bulk T cells
Cd3-
BV605-Cd3
Cd3-
NK Cells
Nkp46-
PE-Cy5-Nkp46
Nkp46-
DCs
Cd11c-
BV570-Cd11c (Itgax)
Cd11c-
Cd11b+ Phagocytes
Cd11b-
BB630-Cd11b (Itgam)
Cd11b-
B Cells
PE-Cd22
Cd11b+
Cd14+
Spark YG570-Cd14
Cd14+
Monocytes
Macrophages
VioGreen-Cd68
Cd68+
M2 Macrophages
M1 Macrophages
BUV737-Cd86
APC-Fire 810-Cd163
Cd68-
M2 Monocytes
M1 Monocytes
Cd11b-
Cd11b+ Iba1+ Phagocytes
Cd11b+ Iba1+ Tmem119+ Phagocytes
Cd11b+ Tmem119+ Phagocytes
PE-Cy7-Tmem119
AF647-Iba1
Cd3-
Gfap-
Gfap+
APC-Gfap
Gfap+
Gfap+ Only Cells
Gfap+ S100b+ Cells
PerCP-S100b
Gfap-
S100b+ Only Cells
PerCP-S100b
Cd3+
Bulk Cd8+ T cells
BV480-Cd8a
Cd3+
Bulk Cd4+ T cells
Pacific Blue-Cd4
Cd4+
TREG
TCON
BV786-Cd127 (Il-7ra)
BV711-Cd25 (Il-2ra)
Cd127-/lo
Cd25-/lo/+/hi
TH2
PerCP-Cy5.5-Cd194 (Ccr4)
Cd194+
TH17
PE-Dazzle594-Cd196 (Ccr6)
TFH
PE-Fire810-Pd-1
BV421-Cd185 (Cxcr5)
TH1
BV650-Cd183 (Cxcr3)
Memory T Cell Gating for All T Cell Subsets
Memory T Cell Gating for All T Cell Subsets
Naïve-Like
TCM
TEF
TEM
BV750-Cd45ra
AF532-Cd62l (L-selectin)

**Supplementary Figure 5: Gating Strategy for Intracellular Median Fluorescent Intensity (MFI) Quantification of Effector Molecules in Brain Single-Cell Suspensions Following Acute *Ex Vivo* Lipopolysaccharide (LPS) Stimulation**

Intracellular effector molecule tracking in central (brain) immune cells following an acute 6-hour *ex vivo* lipopolysaccharide (LPS) stimulation protocol (N = 5 per genotype). Gating logic was confirmed using unstained, unstimulated, single-color, and FMO control samples. Details for the intracellular staining panel are in **Supplementary Table 3**. To visualize the gating workflow, raw FCS files from all biological replicates across all cohorts were concatenated into a single master file (15 files total), (**A.**). Sequential manual gating strategy used to isolate single, live mononuclear cells. (**B.**). Deconvolution of the master dataset back into individual cohorts (WT, GKI, LKO) via SampleID keyword tracking, followed by targeted gating to evaluate expression levels for the following effector molecules: Il-1β, Il-2. Il-4, Il-6, Il-10, Il-13, Il-17, Tnf, Ifnγ, Perforin, Granzyme B, and Gm-csf.

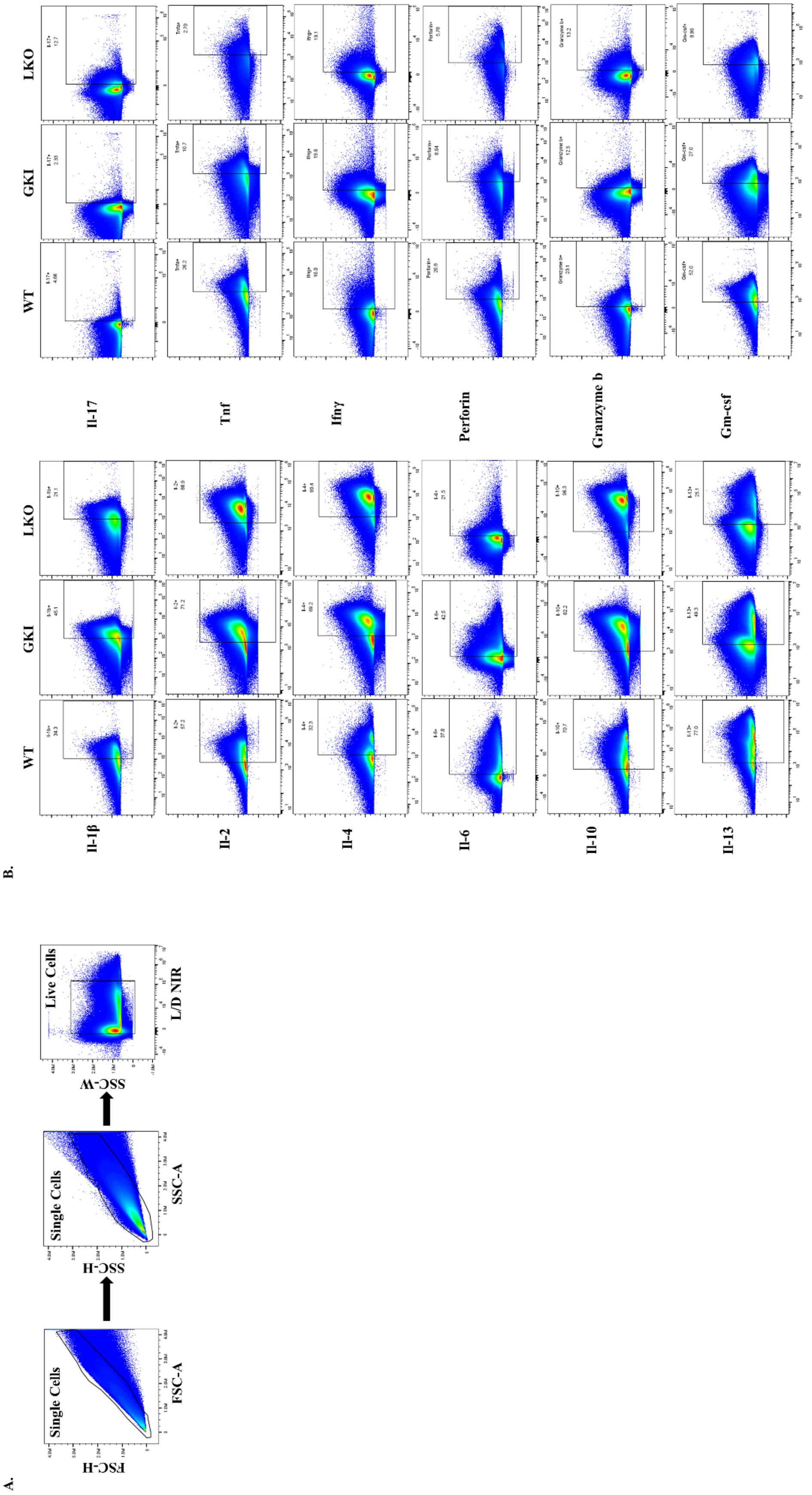
A.
Single Cells
FSC-H
FSC-A
Single Cells
SSC-H
SSC-A
Live Cells
SSC-W
L/D NIR
B.
WT
GKI
LKO
Il-1β
Il-2
Il-4
Il-6
Il-10
Il-13
Il-17
Tnf
Ifnγ
Perforin
Granzyme b
Gm-csf

**Supplementary Figure 6: Gating Strategy for Intracellular Median Fluorescent Intensity (MFI) Quantification of Effector Molecules in Peripheral Blood (PBMCs) and Splenocyte Single-Cell Suspensions Following Acute *Ex Vivo* Lipopolysaccharide (LPS) Stimulation**

Intracellular effector molecule tracking in combined peripheral blood mononuclear cells (PBMCs) and splenocytes following an acute 6-hour *ex vivo* lipopolysaccharide (LPS) stimulation protocol (N = 5 per genotype). Gating logic was confirmed using unstained, unstimulated, single-color, and FMO control samples. Details for the intracellular staining panel are in **Supplementary Table 3**. To visualize the gating workflow, raw FCS files from all biological replicates across all cohorts were concatenated into a single master file (15 files total), (**A.**). Sequential manual gating strategy used to isolate single, live mononuclear cells. (**B.**). Deconvolution of the master dataset back into individual cohorts (WT, GKI, LKO) via SampleID keyword tracking, followed by targeted gating to evaluate expression levels for the following effector molecules: Il-1β, Il-2. Il-4, Il-6, Il-10, Il-13, Il-17, Tnf, Ifnγ, Perforin, Granzyme B, and Gm-csf.

A.

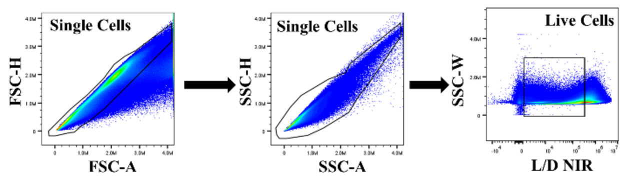
Single Cells
FSC-H
FSC-A
Single Cells
SSC-H
SSC-A
Live Cells
SSC-W
L/D NIR

B.

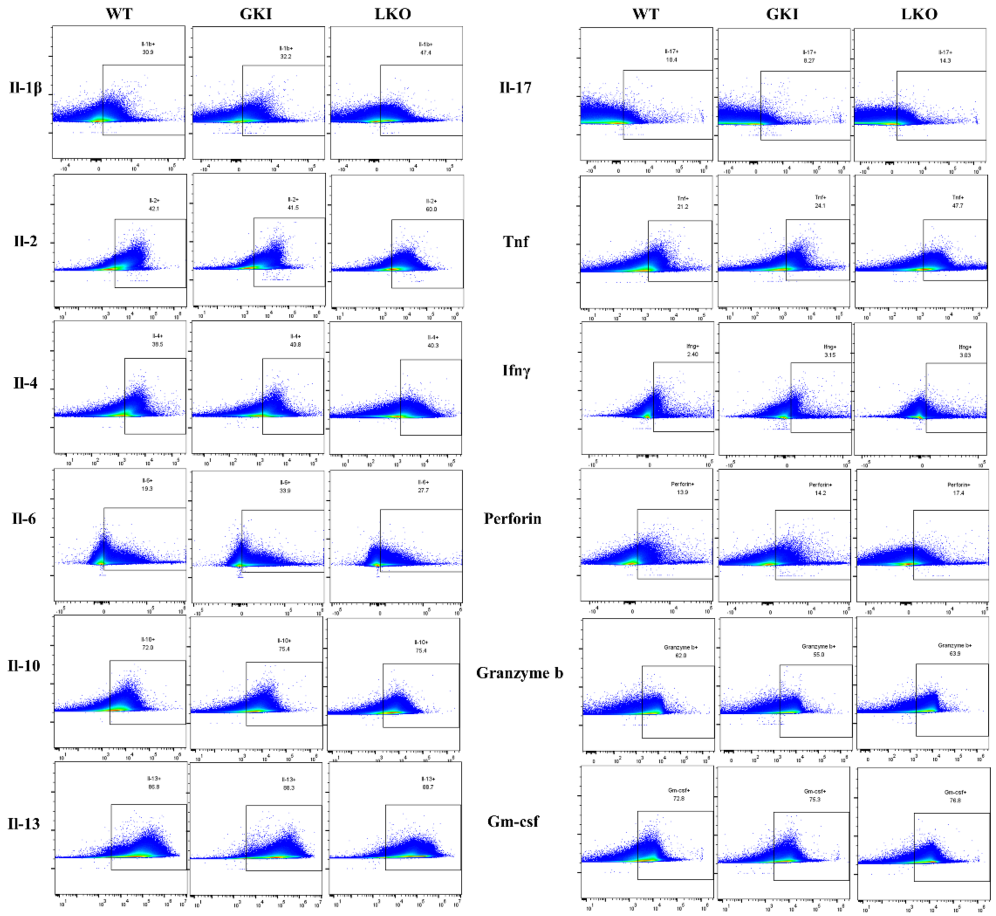
WT
GKI
LKO
Il-1β
Il-2
Il-4
Il-6
Il-10
Il-13
WT
GKI
LKO
Il-17
Tnf
Ifnγ
Perforin
Granzyme b
Gm-csf

**Supplementary Figure 7: Example of Workflow for Semi-Automated Computational Pipeline for High-Dimensional tSNE Clustering and FlowSOM Meta-Cluster Mapping**

Schematic workflow illustrating unbiased topologic discovery across (**A.**) central (brain) and (**B.**) peripheral (PBMCs/splenocytes) immune compartments (N = 5 per genotype). Individual FlowJo™ FCS files were downsampled using the DownSample plugin (BD Life Sciences©, V3) to a fixed depth of 1,000,000 total events, resulting in final concatenated populations of 639,048 (brain) and 665,503 (periphery) per tissue system to normalize processing weights. Downsampled files were concatenated and evaluated using the tSNE analysis plugin in FlowJo™ under optimized hyperparameters: iterations = 1000, perplexity = 30, KNN algorithm = Exact, Gradient Algorithm = Barnes-Hut. Unsupervised population boundaries were resolved using X-Shift plugin (BD Life Sciences©, v.1.4.1) to mathematically determine absolute cluster counts based on an angular distance metric (k = 37). Fine-resolution meta-clustering was executed via the FlowSOM plugin (BD Life Sciences©, v3.0.18), resolving 15 meta-clusters in the central immune system dataset and 28 meta-clusters in the peripheral immune system dataset. Individual cluster phenotypes were parsed using the Cluster Explorer plugin, where a cluster was assigned an identity category if its relative fluorescence expression matched or exceeded a strict threshold of ≥ $10^4$ MFI. Backbone tracking markers included: Cd8$^+$ (Cd8$^+$ T cells), Cd4$^+$ (Cd4$^+$ T cells), Cd11c$^+$ (DCs), Nkp46$^+$ (NK cells), Cd22$^+$ (B cells), Cd11b$^+$ (Cd11b$^+$ phagocytes), Tmem119$^+$ (Tmem119$^+$ phagocytes), Iba1$^+$ (Iba1$^+$ phagocytes), Gfap$^+$ (Gfap$^+$ astrocytes), and S100b$^+$ (S100b$^+$ astrocytes). Clusters failing to exceed this expression threshold across all core lineage parameters were excluded from the final presentation, yielding a refined cluster count of 12 distinct central immune clusters and 15 peripheral immune clusters. Secondary phenotypic profiles (defined by other markers in **Supplementary Table 2**) meeting the expression threshold are indicated in parentheses. Population frequencies were subsequently deconvoluted by genotype via SampleID tags for comparative analysis.

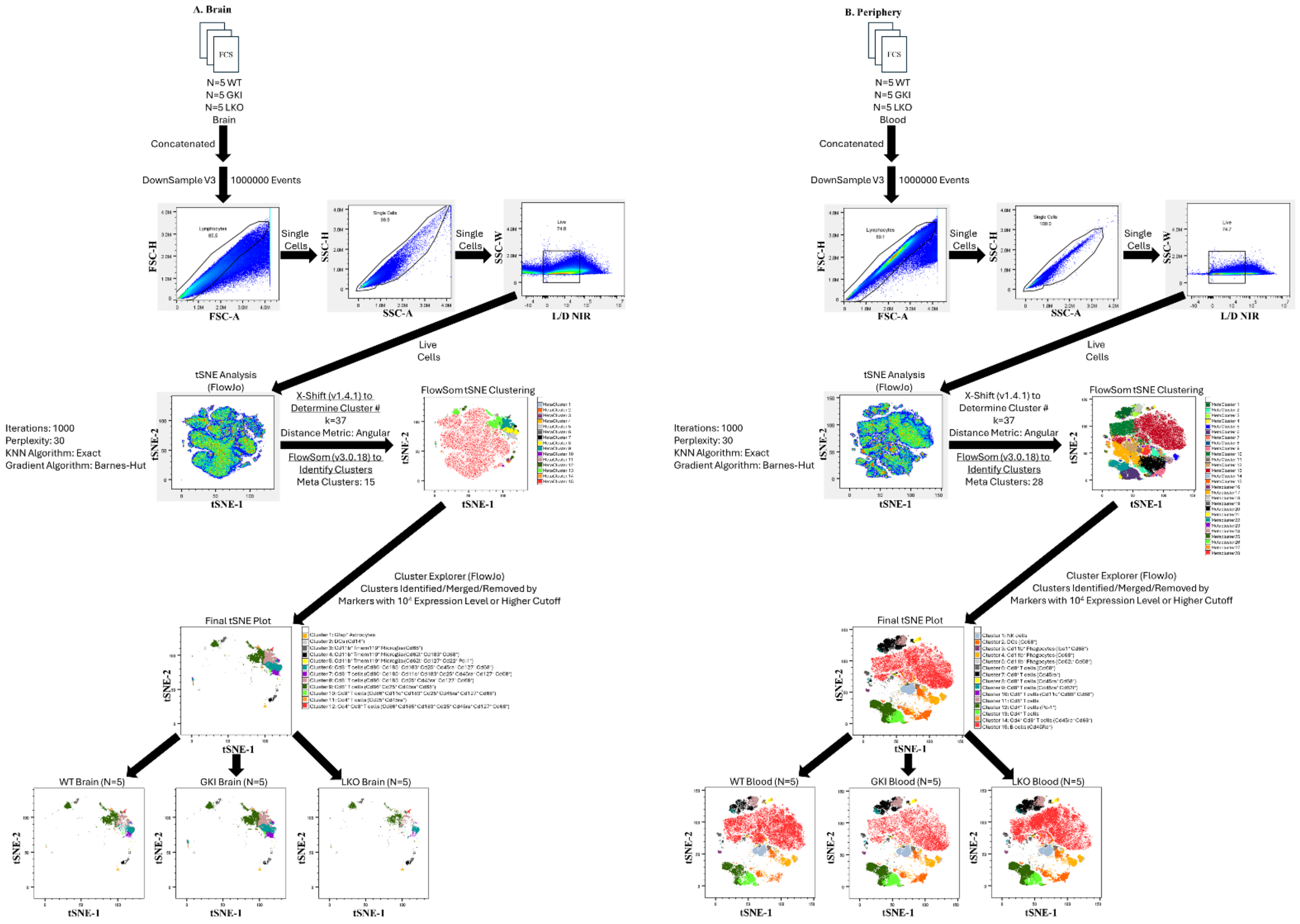
A. Brain
FCS
N=5 WT
N=5 GKI
N=5 LKO
Brain
Concatenated
DownSample V3
1000000 Events
FSC-H
FSC-A
Single Cells
SSC-H
SSC-A
Single Cells
SSC-W
L/D NIR
Live Cells
tSNE Analysis (FlowJo)
Iterations: 1000
Perplexity: 30
KNN Algorithm: Exact
Gradient Algorithm: Barnes-Hut
X-Shift (v1.4.1) to Determine Cluster #
k=37
Distance Metric: Angular
FlowSom (v3.0.18) to Identify Clusters
Meta Clusters: 15
FlowSom tSNE Clustering
tSNE-1
tSNE-2
Cluster Explorer (FlowJo)
Clusters Identified/Merged/Removed by
Markers with 10⁴ Expression Level or Higher Cutoff
Final tSNE Plot
WT Brain (N=5)
GKI Brain (N=5)
LKO Brain (N=5)
B. Periphery
FCS
N=5 WT
N=5 GKI
N=5 LKO
Blood
Concatenated
DownSample V3
1000000 Events
Single Cells
Single Cells
Live Cells
tSNE Analysis (FlowJo)
Iterations: 1000
Perplexity: 30
KNN Algorithm: Exact
Gradient Algorithm: Barnes-Hut
X-Shift (v1.4.1) to Determine Cluster #
k=37
Distance Metric: Angular
FlowSom (v3.0.18) to Identify Clusters
Meta Clusters: 28
FlowSom tSNE Clustering
Cluster Explorer (FlowJo)
Clusters Identified/Merged/Removed by
Markers with 10⁴ Expression Level or Higher Cutoff
Final tSNE Plot
WT Blood (N=5)
GKI Blood (N=5)
LKO Blood (N=5)

**Supplementary Figure 8: Volcano Plot Screen of Additional Immune Cell Subset Shifts in GKI and LKO Models Under Basal Conditions**

Statistical screening of additional homeostatic immune subset frequencies across (**A.**) central (brain) and (**B.**) peripheral (PBMCs/splenocytes) immune systems (N = 5 per genotype). Gated frequencies were calculated as a percentage of total parent events: Live Cell % = (Gated Subset Cell Count/Total Live Cells) x 100. Fold-change (FC) boundaries were calculated for GKI/WT and LKO/WT configurations. Group averages were transformed into Log2 Fold Change (Log2 FC) and plotted against their corresponding statistical weight, expressed as -Log10 *P-value*.  Parametric datasets were analyzed using a Brown-Forsythe and Welch ANOVA with Dunnet's T3 correction test and non-parametric datasets were evaluated via a Kruskal-Wallis test with a Dunn's multiple comparison test. Phenotypes with *P-value* ≤ 0.05 are highlighted in red for positive fold change and blue for negative fold change along with the points being labeled with their descriptive lineage names.

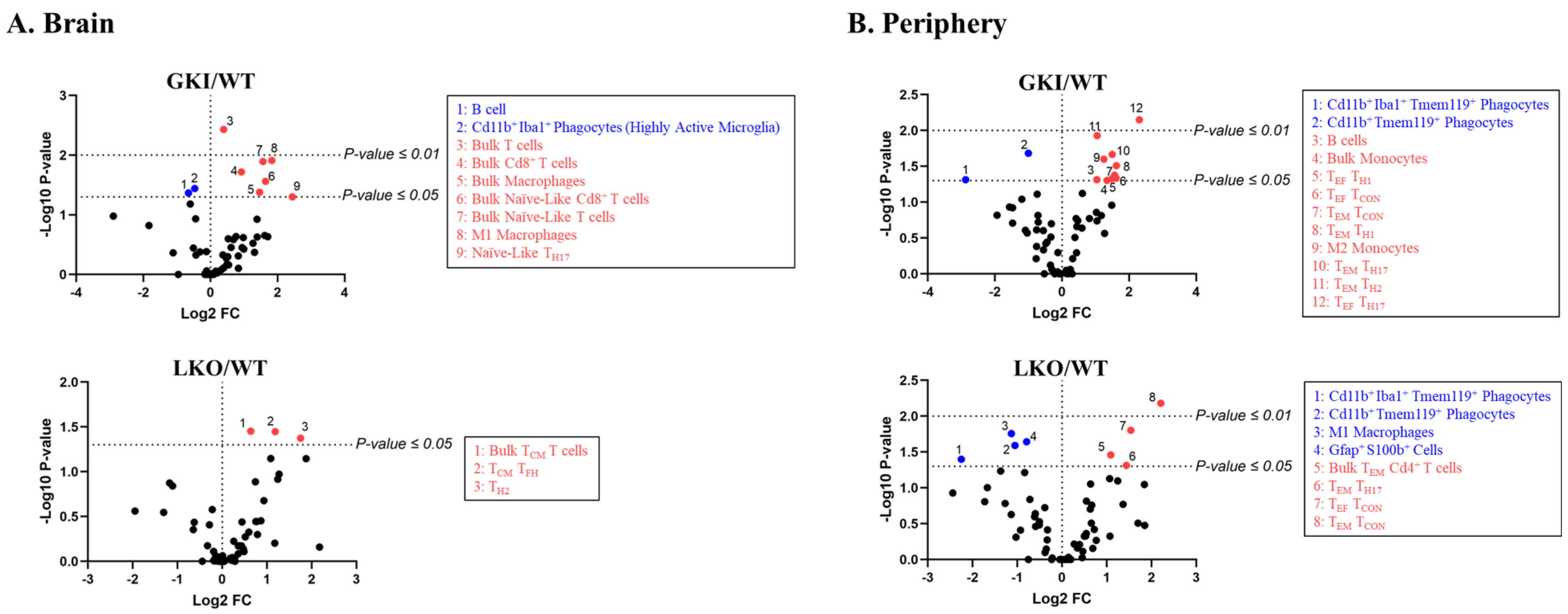
A. Brain
GKI/WT
-Log10 P-value
Log2 FC
P-value ≤ 0.01
P-value ≤ 0.05
1: B cell
2: Cd11b+Iba1+ Phagocytes (Highly Active Microglia)
3: Bulk T cells
4: Bulk Cd8+ T cells
5: Bulk Macrophages
6: Bulk Naïve-Like Cd8+ T cells
7: Bulk Naïve-Like T cells
8: M1 Macrophages
9: Naïve-Like TH17
LKO/WT
P-value ≤ 0.05
1: Bulk TCM T cells
2: TCM TFH
3: TH2
B. Periphery
GKI/WT
P-value ≤ 0.01
P-value ≤ 0.05
1: Cd11b+Iba1+ Tmem119+ Phagocytes
2: Cd11b+Tmem119+ Phagocytes
3: B cells
4: Bulk Monocytes
5: TEF TH1
6: TEF TCON
7: TEM TCON
8: TEM TH1
9: M2 Monocytes
10: TEM TH17
11: TEM TH2
12: TEF TH17
LKO/WT
P-value ≤ 0.01
P-value ≤ 0.05
1: Cd11b+Iba1+ Tmem119+ Phagocytes
2: Cd11b+Tmem119+ Phagocytes
3: M1 Macrophages
4: Gfap+S100b+ Cells
5: Bulk TEM Cd4+ T cells
6: TEM TH17
7: TEF TCON
8: TEM TCON

**Supplementary Figure 9: Manual Flow Cytometry Validation Gating Trees for Significant Brain T cell Populations**

Detailed gating paths for the specific central (brain) immune system T cell lineages identified as significantly altered in **Figure 4** (N = 5 per genotype). Complete antibody specifications are provided in **Supplementary Table 2**. Individual replicates were concatenated (15 files total) to build standard gating pathways, followed by SampleID deconvolution. The panel displays explicit sequential gating tracks used to isolate and quantify: bulk T cells (Cd3$^{+}$), bulk memory T cells (Cd3$^{+}$Cd45ra$^{+/-}$Cd62l$^{+/-}$), bulk Cd8$^{+}$ T cells (Cd3$^{+}$Cd8$^{+}$), bulk memory Cd8$^{+}$ T cells (Cd3$^{+}$Cd8$^{+}$Cd45ra$^{+/-}$Cd62l$^{+/-}$), memory T$_{H17}$ (Cd3$^{+}$ Cd4$^{+}$ Cd25$^{-/lo/+/hi}$ Cd127$^{+/hi}$ Cd194$^{+}$ Cd196$^{+}$ Cd62l$^{+/-}$ Cd45ra$^{+/-}$), T$_{H2}$ (Cd3$^{+}$ Cd4$^{+}$ Cd25$^{-/lo/+/hi}$ Cd127$^{+/hi}$ Cd194$^{+}$), and memory T$_{FH}$ (Cd3$^{+}$ Cd4$^{+}$ Cd25$^{-/lo/+/hi}$ Cd127$^{+/hi}$ Cd185$^{+}$ Pd-1$^{+}$ Cd62l$^{+/-}$ Cd45ra$^{+/-}$).

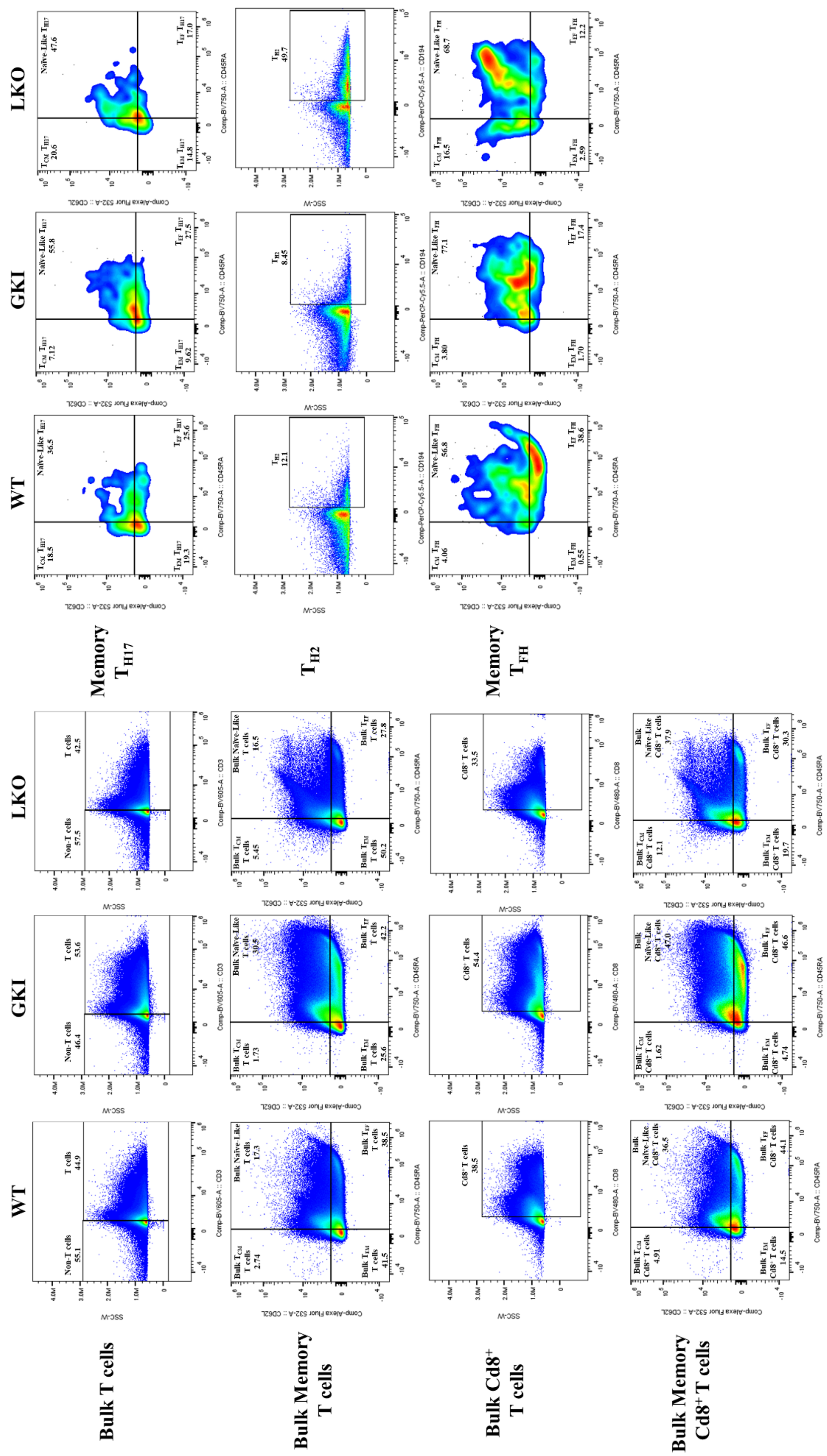
WT
GKI
LKO
Bulk T cells
Bulk Memory T cells
Bulk Cd8+ T cells
Bulk Memory Cd8+ T cells
Memory TH17
TH2
Memory TFH

**Supplementary Figure 10: Manual Flow Cytometry Gating Path for the Isolation of Tissue Stem Cell Memory T cells ($T_{SCM}$)**

Technical verification of the gating cascade used to track central tissue-resident stem cell memory subsets (N = 5 per genotype). Antibody data in **Supplementary Table 2** and lineage definitions, gating hierarchies, and peer-reviewed classification references for each identified subset are detailed in **Supplementary Table 4**. Following live single-cell isolation on the 15 file concatenated master template, events were branched into bulk $Cd3^+$ and bulk $Cd3^+Cd8^+$ T cells. Within these parental gates, high defined $T_{SCM}$ T cells were determined by $Cd45ra^+Cd62l^+Cd183(Cxcr3)^+$ gating.

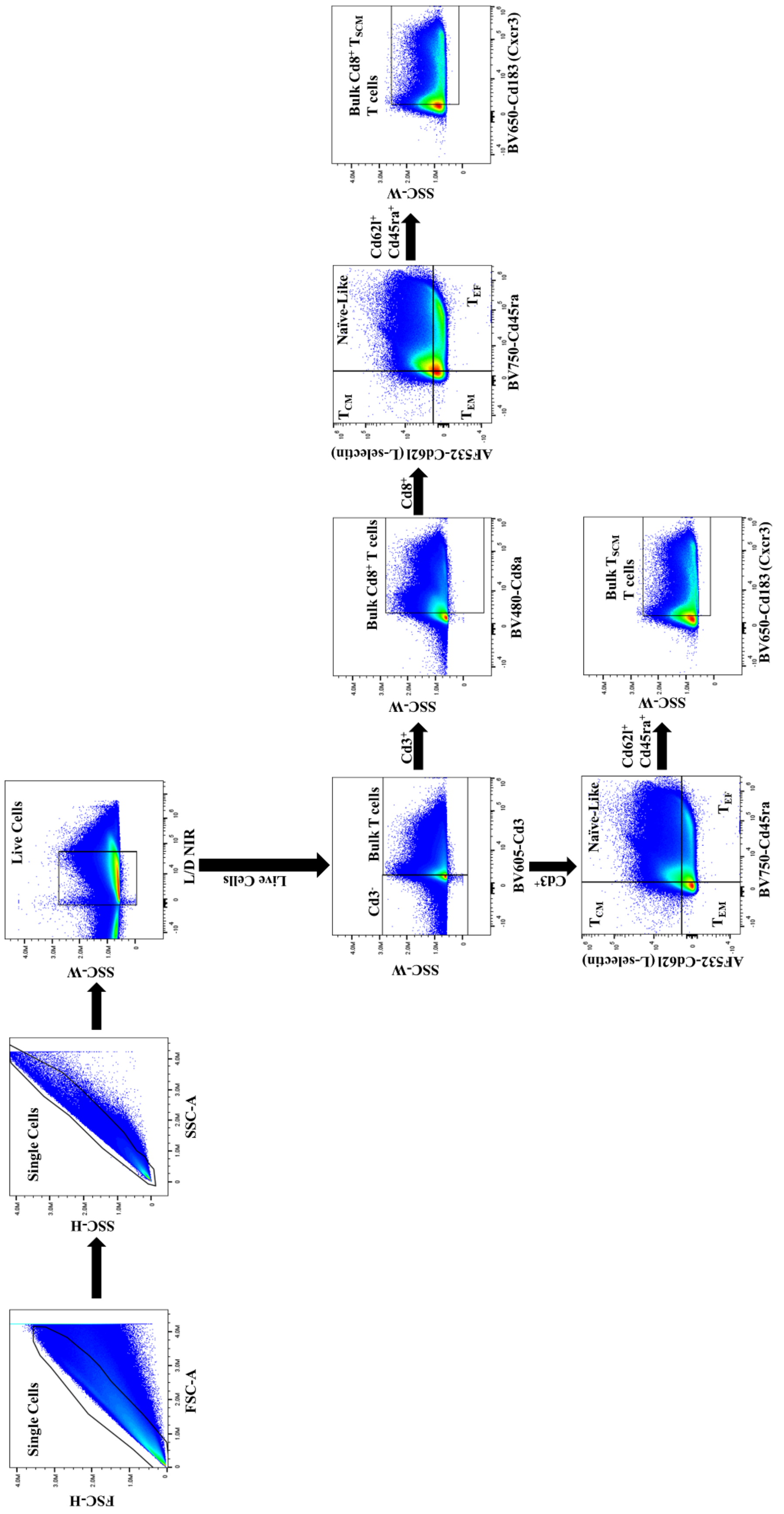

Single Cells
FSC-H
FSC-A
Single Cells
SSC-H
SSC-A
Live Cells
SSC-W
L/D NIR
Live Cells
Bulk T cells
Cd3-
SSC-W
BV605-Cd3
Cd3+
Bulk Cd8+ T cells
SSC-W
BV480-Cd8a
Cd8+
T_CM
Naïve-Like
T_EM
T_EF
AF532-Cd62l (L-selectin)
BV750-Cd45ra
Cd62l+ Cd45ra+
Bulk Cd8+ T_SCM T cells
SSC-W
BV650-Cd183 (Cxcr3)
T_CM
Naïve-Like
T_EM
T_EF
AF532-Cd62l (L-selectin)
BV750-Cd45ra
Cd62l+ Cd45ra+
Bulk T_SCM T cells
SSC-W
BV650-Cd183 (Cxcr3)

**Supplementary Figure 11: Manual Flow Cytometry Validation Gating Trees for Significant Peripheral (PBMCs/Splenocytes) T cell Populations**

Detailed gating paths for the specific peripheral (PBMCs/Splenocytes) immune system T cell lineages identified as significantly altered in **Figure 4** (N = 5 per genotype). Complete antibody specifications are provided in **Supplementary Table 2**. Individual replicates were concatenated (15 files total) to build standard gating pathways, followed by SampleID deconvolution. The panel displays explicit sequential gating tracks used to isolate and quantify: bulk memory $Cd4^{+}$ T cells ($Cd3^{+}Cd4^{+}Cd62l^{+/-}Cd45ra^{+/-}$), memory $T_{CON}$ ($Cd3^{+}$ $Cd4^{+}$ $Cd25^{-/lo/+/hi}$ $Cd127^{+/hi}$ $Cd62l^{+/-}$ $Cd45ra^{+/-}$), memory $T_{H1}$ ($Cd3^{+}$ $Cd4^{+}$ $Cd25^{-/lo/+/hi}$ $Cd127^{+/hi}$ $Cd183^{+}$ $Cd62l^{+/-}$ $Cd45ra^{+/-}$), memory $T_{H2}$ ($Cd3^{+}$ $Cd4^{+}$ $Cd25^{-/lo/+/hi}$ $Cd127^{+/hi}$ $Cd194^{+}$ $Cd62l^{+/-}$ $Cd45ra^{+/-}$), and memory $T_{H17}$ ($Cd3^{+}$ $Cd4^{+}$ $Cd25^{-/lo/+/hi}$ $Cd127^{+/hi}$ $Cd194^{+}$ $Cd196^{+}$ $Cd62l^{+/-}$ $Cd45ra^{+/-}$).

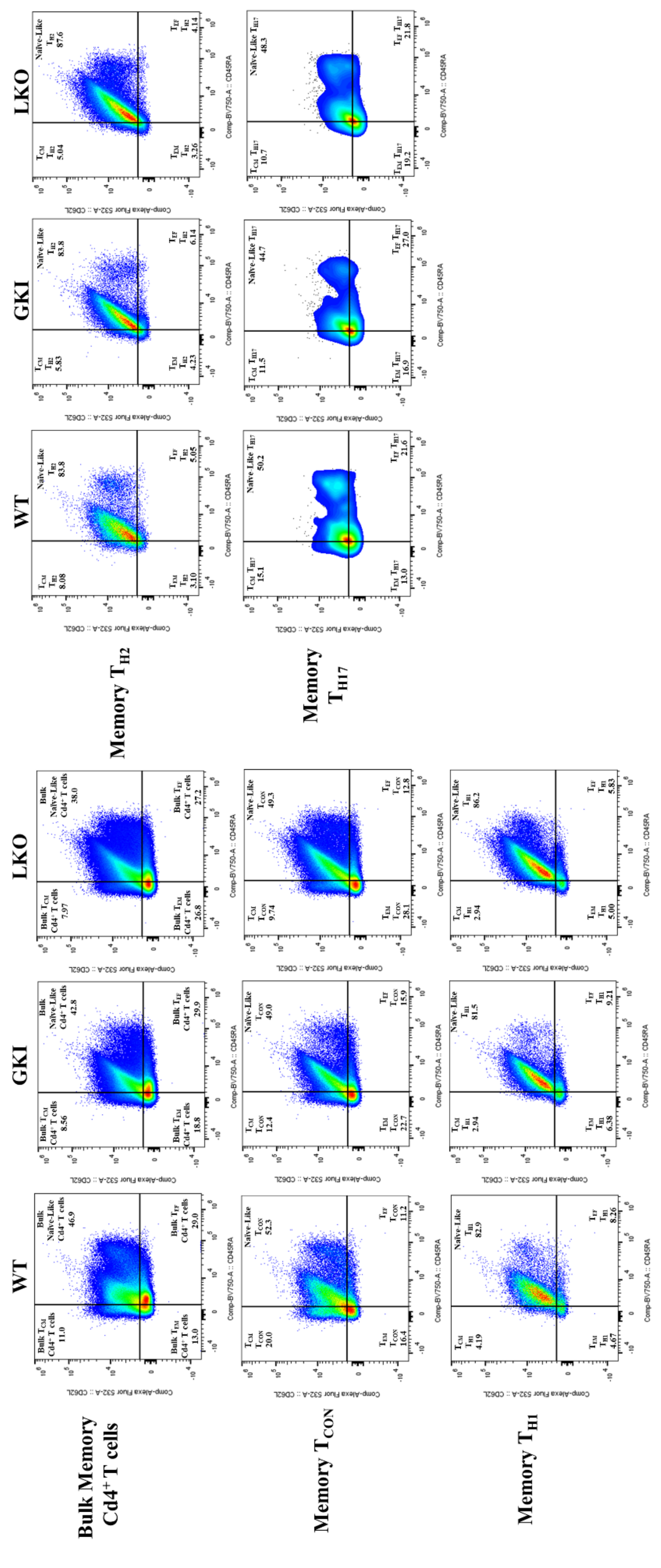
WT
GKI
LKO
Memory TH2
Memory TH17
WT
GKI
LKO
Bulk Memory Cd4+ T cells
Memory TCON
Memory TH1

**Supplementary Figure 12: Volcano Plot Screen of Additional Intracellular Effector Molecule Shifts in GKI and LKO Models Following Acute *Ex Vivo* Lipopolysaccharide (LPS) Stimulation**

Statistical screening of additional effector molecular producing immune cell subset frequencies across (**A.**) central (brain) and (**B.**) peripheral (PBMCs/splenocytes) immune cells following a 6-hour *ex vivo* lipopolysaccharide (LPS) challenge (N = 5 per genotype). Comparisons between fold change of the integrated MFI (iMFI = [nMFI reading of effector molecule] * [percentage positive for marker within that immune cell subset]) for each effector molecule expression was found between models. Fold-change (FC) boundaries were calculated for GKI/WT and LKO/WT configurations. Group averages were transformed into Log2 Fold Change (Log2 FC) and plotted against their corresponding statistical weight, expressed as -Log10 *P-value*. Parametric datasets were analyzed using a Brown-Forsythe and Welch ANOVA with Dunnet's T3 correction test and non-parametric datasets were evaluated via a Kruskal-Wallis test with a Dunn's multiple comparison test. Phenotypes with *P-value* $\leq 0.05$ are highlighted in red for positive fold change and blue for negative fold change along with the points being labeled with their descriptive lineage names.

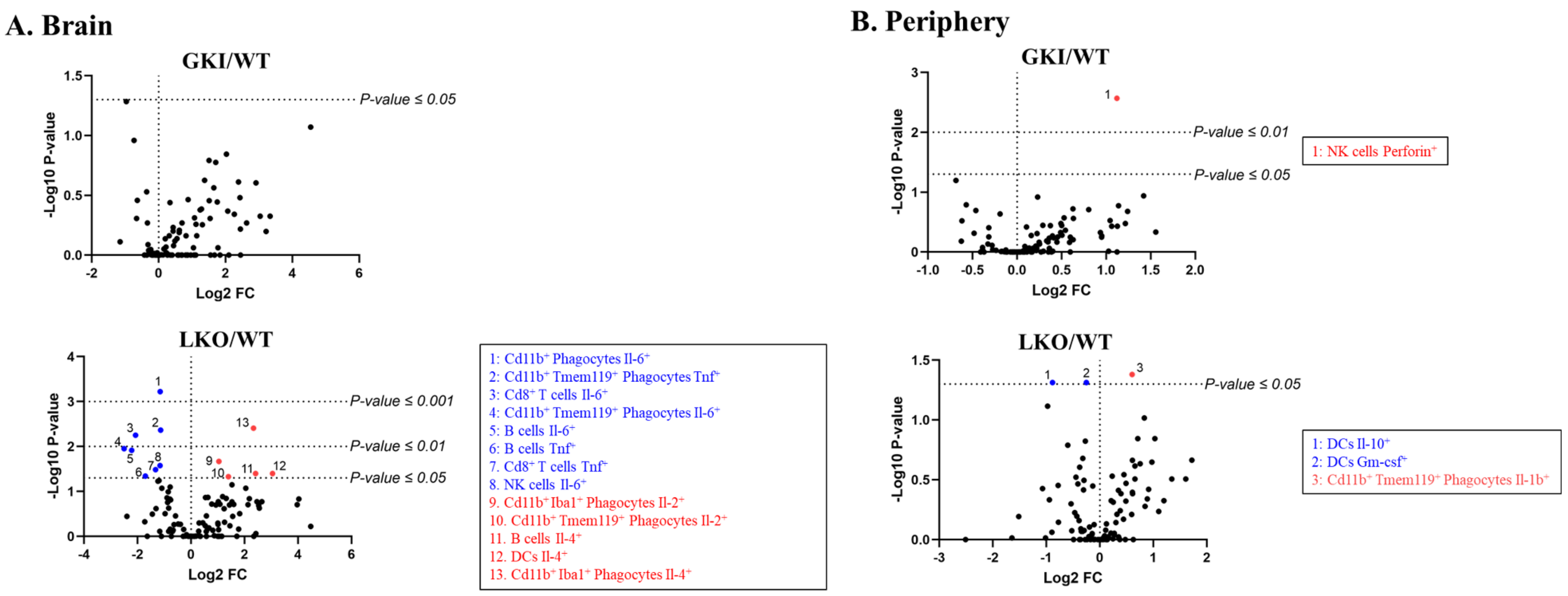
A. Brain
GKI/WT
-Log10 P-value
Log2 FC
P-value ≤ 0.05
LKO/WT
P-value ≤ 0.001
P-value ≤ 0.01
P-value ≤ 0.05
1: Cd11b+ Phagocytes Il-6+
2: Cd11b+ Tmem119+ Phagocytes Tnf+
3: Cd8+ T cells Il-6+
4: Cd11b+ Tmem119+ Phagocytes Il-6+
5: B cells Il-6+
6: B cells Tnf+
7. Cd8+ T cells Tnf+
8. NK cells Il-6+
9. Cd11b+ Iba1+ Phagocytes Il-2+
10. Cd11b+ Tmem119+ Phagocytes Il-2+
11. B cells Il-4+
12. DCs Il-4+
13. Cd11b+ Iba1+ Phagocytes Il-4+
B. Periphery
GKI/WT
P-value ≤ 0.01
P-value ≤ 0.05
1: NK cells Perforin+
LKO/WT
P-value ≤ 0.05
1: DCs Il-10+
2: DCs Gm-csf+
3: Cd11b+ Tmem119+ Phagocytes Il-1b+

**Supplemental Figure 13: Quantitative PCR (qPCR) Profiling of *Pdcd1* and *Lag3* in Central and Peripheral Immune Cells**

Relative mRNA expression levels of *Pdcd1* (programmed cell death 1 and *Lag3* (lymphocyte activation gene 3) determined via qPCR in (**A.**) central (brain) and (**B.**) peripheral (PBMCs/splenocytes) across WT, GKI, and LKO cohorts (N = 5 per genotypes). qPCR primers can be found in **Supplementary Table 1**. Relative gene expression ($2^{-\Delta\Delta CT}$) was calculated for each individual biological replicate, and bar graphs were made (mean ± SEM). Exact *P-values* are annotated above brackets. Statistical analysis was done using a Brown-Forsythe and Welch ANOVA with a Dunnett's T3 multiple comparisons test. A Benjamini-Hochberg FDR correction was applied for all *p-values* across all comparisons.

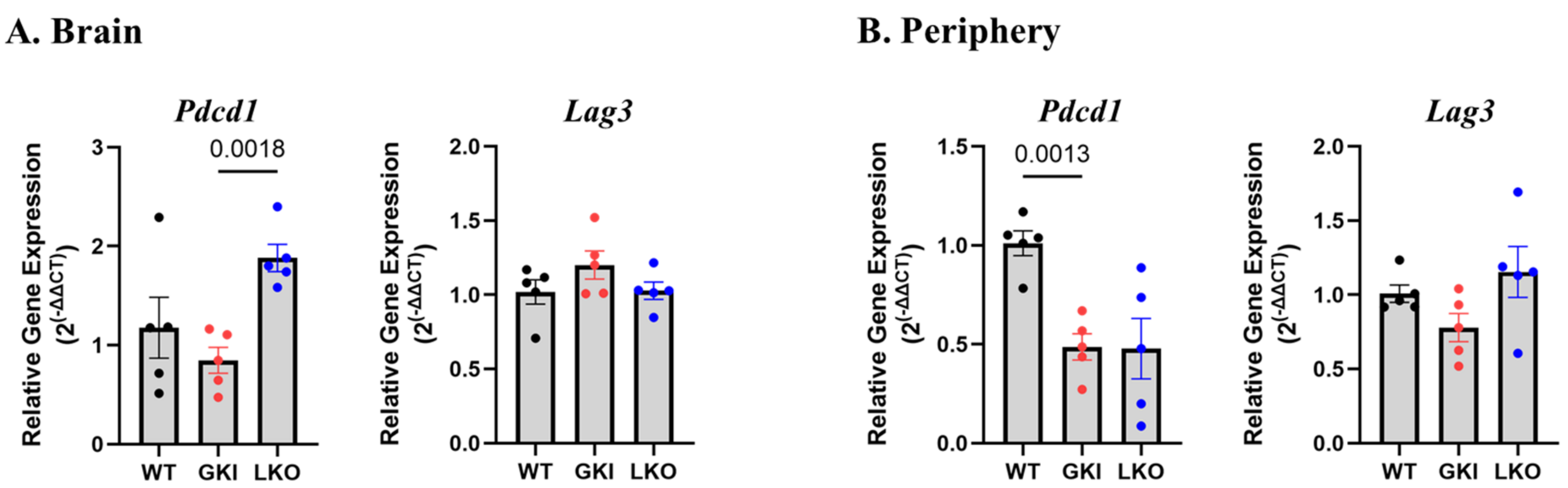

A. Brain
B. Periphery
Pdcd1
Lag3
Pdcd1
Lag3
0.0018
0.0013
Relative Gene Expression ($2^{(-\Delta\Delta CT)}$)
WT
GKI
LKO

**Supplemental Figure 14: Flow Cytometry Gating for Targeted Pd-1 Expression Profiling Across Adaptive and Innate Immune Lineages**

Detailed gating paths for the specific (**A.**) central (brain) and (**B.**) peripheral (PBMCs/splenocytes) immune system T cell lineages identified as significantly altered in **Figure 9** (N = 5 per genotype). Complete antibody specifications are provided in **Supplementary Table 2**. Individual replicates were concatenated (15 files total) to build standard gating pathways, followed by SampleID deconvolution. The panel displays explicit sequential gating tracks used to isolate and quantify Pd-1 expression: Cd4$^{+}$ T cells, Cd8$^{+}$ T cells, $T_{REG}$ cells, $T_{H1}$ cells, $T_{H2}$ cells, $T_{H17}$ cells, B cells, NK cells, DCs, and Cd11b$^{+}$ phagocytes.

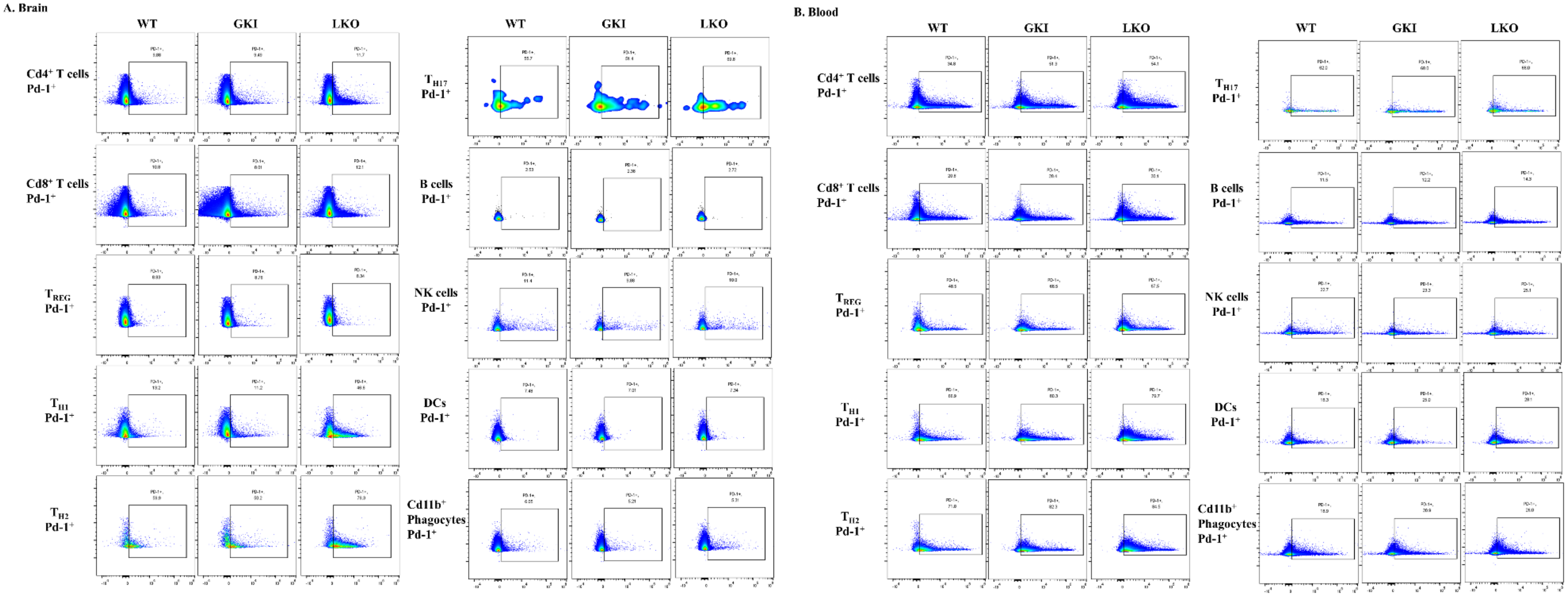
A. Brain
WT
GKI
LKO
Cd4⁺ T cells Pd-1⁺
Cd8⁺ T cells Pd-1⁺
T_REG Pd-1⁺
T_H1 Pd-1⁺
T_H2 Pd-1⁺
T_H17 Pd-1⁺
B cells Pd-1⁺
NK cells Pd-1⁺
DCs Pd-1⁺
Cd11b⁺ Phagocytes Pd-1⁺
B. Blood
WT
GKI
LKO
Cd4⁺ T cells Pd-1⁺
Cd8⁺ T cells Pd-1⁺
T_REG Pd-1⁺
T_H1 Pd-1⁺
T_H2 Pd-1⁺
T_H17 Pd-1⁺
B cells Pd-1⁺
NK cells Pd-1⁺
DCs Pd-1⁺
Cd11b⁺ Phagocytes Pd-1⁺

**Supplementary Figure 15: Integrated Median Fluorescence Intensity (iMFI) Fold Changes of Pd-1 Expression Across Myeloid and Innate Lymphoid Lineages**

Quantitative assessment of integrated median fluorescence intensity (iMFI = [nMFI reading of effector molecule] * [percentage positive for marker within that immune cell subset]) fold changes for surface Pd-1 expression under basal homeostatic conditions in (**A.**) central (brain) and (**B.**) peripheral (PBMCs/splenocytes) immune cells across WT, GKI, and LKO cohorts (N = 5 per genotype). Bar graphs plot individual biological replicates tracking iMFI of Pd-1 (mean ± SEM). Exact *P-values* are annotated above brackets. For parametric data, statistical analysis was done using a Brown-Forsythe and Welch ANOVA with a Dunnett's T3 multiple comparisons test. For nonparametric data, a Kruskal-Wallis test with a Dunn's multiple comparison test was utilized.

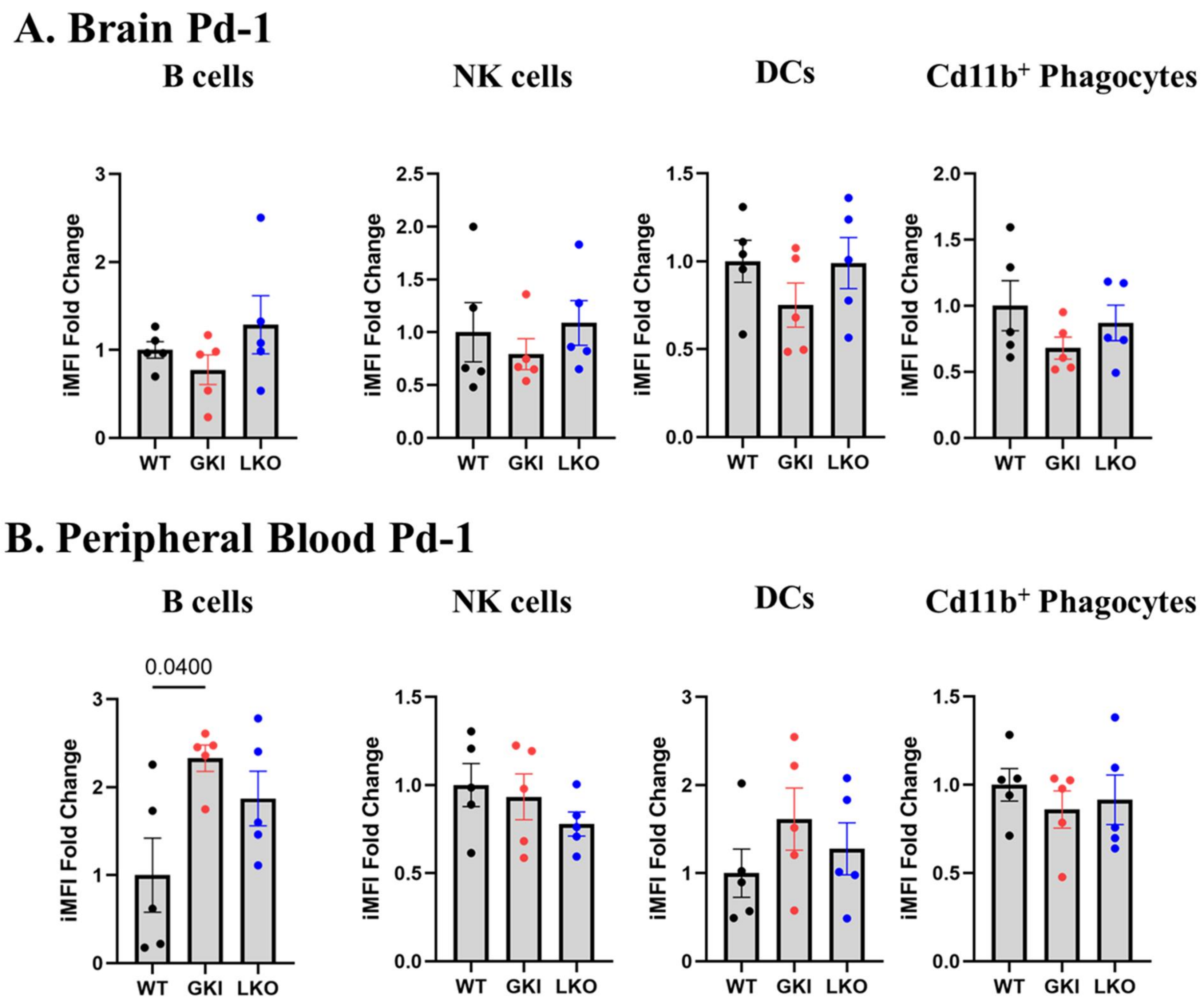

A. Brain Pd-1
B cells
NK cells
DCs
Cd11b+ Phagocytes
iMFI Fold Change
WT
GKI
LKO
B. Peripheral Blood Pd-1
B cells
NK cells
DCs
Cd11b+ Phagocytes
0.0400
iMFI Fold Change
WT
GKI
LKO

### SUPPLEMENTARY TABLES

**Table S1. All TaqMan™ Gene Expression qPCR Assays Used from Thermo Fisher Scientific©**

| Target | Assay ID |
|---|---|
| *Aif1 (Iba1)* | Mm00479862_g1 |
| *Cd22* | Mm00515432_m1 |
| *Cd3e* | Mm01179194_m1 |
| *Cd4* | Mm00442754_m1 |
| *Cd8a* | Mm01182107_g1 |
| *Csf2* | Mm01290062_m1 |
| *Gfap* | Mm01253033_m1 |
| *Gzmb* | Mm00442837_m1 |
| *Ifng* | Mm01168134_m1 |
| *Il10* | Mm01288386_m1 |
| *Il13* | Mm00434204_m1 |
| *Il17a* | Mm00439618_m1 |
| *Il1b* | Mm00434228_m1 |
| *Il2* | Mm00434256_m1 |
| *Il4* | Mm00445259_m1 |
| *Il6* | Mm00446190_m1 |
| *Itgam (Cd11b)* | Mm00434455_m1 |

| | |
|---|---|
| *Itgax (Cd11c)* | Mm00498701_m1 |
| *Lag3* | Mm00493071_m1 |
| *Ncr1 (Nkp46)* | Mm01337324_g1 |
| *Pdcd1 (Pd-1)* | Mm01285676_m1 |
| *Prf1* | Mm00812512_m1 |
| *S100b* | Mm00485897_m1 |
| *Tmem119* | Mm00525305_m1 |
| *Tnf* | Mm00443258_m1 |

| Table S2. All Antibodies Used for Immunophenotyping Flow Cytometry Panel | | | | | |
|---|---|---|---|---|---|
| **Marker** | **Intracellular or Extracellular?** | **Clone** | **Fluorophore** | **Channel Examined on Cytek® Aurora** | **Company** |
| Live/Dead™ Fixable Near-IR Dead Cell Stain | Intra | N/A | Near-IR | R7 | Invitrogen™ |
| Cd3 | Extra | 17A2 | BV605™ | V10 | Biolegend® |
| Cd8a | Extra | 53-6.7 | BV480™ | V5 | BD Biosciences© |
| Cd4 | Extra | GK1.5 | Pacific Blue (PB)™ | V3 | Biolegend® |
| Cd127 (Il-7ra) | Extra | SB/199 | BV786™ | V15 | BD Biosciences© |
| Cd25 (Il-2ra) | Extra | PC61 | BV711™ | V13 | BD Biosciences© |
| Pd-1 | Extra | 29F.1A12 | PE-Fire™ 810 | YG10 | Biolegend® |
| Cd185 (Cxcr5) | Extra | L138D7 | BV421™ | V1 | Biolegend® |
| Cd183 (Cxcr3) | Extra | CXCR3-173 | BV650™ | V11 | Biolegend® |
| Cd194 (Ccr4) | Extra | 2G12 | PerCP-Cy5.5 | B9 | Biolegend® |

| | | | | | |
|---|---|---|---|---|---|
| Cd196 (Ccr6) | Extra | 29-2L17 | PE-Dazzle™ 594 | YG3 | Biolegend® |
| Cd45ra | Extra | 14.8 | BV750™ | V14 | BD Biosciences© |
| Cd62l (L-selectin) | Extra | MEL-14 | NovaFluor™ Blue 585 | B5 | Invitrogen™; eBioscience™ |
| Cd11c (Itgax) | Extra | N418 | BV570™ | V8 | Biolegend® |
| Nkp46 (Ncr1) | Extra | 29A1.4 | PE-Cy5 | YG5 | Biolegend® |
| Cd22 | Extra | OX-97 | PE | YG1 | Biolegend® |
| Cd11b (Itgam) | Extra | M1/70 | NovaFluor™ Blue 610/30S | B6 | Invitrogen™; eBioscience™ |
| Tmem119 | Extra | V3RT1GOsz | PE-Cy7 | YG9 | Invitrogen™; eBioscience™ |
| Iba-1 (Aif1) | Intra | 20A12.1 | AF647™ | R2 | Millipore Sigma© |
| Cd14 | Extra | M14-23 | Spark YG™ 593 | YG2 | Biolegend® |
| Cd68 | Intra | REA835 | VioGreen™ | V7 | Miltenyi Biotec© |
| Cd86 (B7-2) | Extra | GL1 | BUV737™ | R5 | BD Biosciences© |
| Cd163 | Extra | S15049I | APC-Fire™ 810 | R8 | Biolegend® |

| | | | | | |
|---|---|---|---|---|---|
| Gfap | Intra | REA335 | APC | R1 | Miltenyi Biotec© |
| S100b | Intra | 15F9NB | PerCP | B8 | Novus Biologicals© |

**Table S3. All Antibodies Used for Intracellular Cytokine Production Flow Cytometry Panel**

| Marker | Intracellular or Extracellular? | Clone | Fluorophore | Channel Examined on Cytek® Aurora | Company |
|---|---|---|---|---|---|
| Live/Dead™ Fixable Near-IR Dead Cell Stain | Intra | N/A | Near-IR | R7 | Invitrogen™ |
| Cd3 | Extra | 17A2 | BV570™ | V8 | Biolegend® |
| Cd8a | Extra | 53-6.7 | NovaFluor™ Blue 585 | B5 | Invitrogen™; eBioscience™ |
| Cd4 | Extra | GK1.5 | APC-Cy5.5 | R3 | Southern Biotechnology Associates™ |
| Cd11c (Itgax) | Extra | N418 | NovaFluor Blue 610/30S | B6 | Invitrogen™; eBioscience™ |
| Nkp46 (Ncr1) | Extra | 29A1.4 | BV480™ | V5 | BD Biosciences© |
| Cd22 | Extra | CD22.2 | BV650™ | V10 | BD Biosciences© |
| Cd11b (Itgam) | Extra | M1/70 | SuperBright™ 780 | V15 | Invitrogen™; eBioscience™ |

| | | | | | |
|---|---|---|---|---|---|
| Tmem119 | Extra | V3RT1GOsz | PE-Cy7 | YG9 | Invitrogen™; eBioscience™ |
| Iba-1 (Aif1) | Intra | E4O4W | PE | YG1 | Cell Signaling Technology© |
| Il-1b | Intra | 001 | AF594™ | YG3 | Novus Biologicals© |
| Il-2 | Intra | JES6-5H4 | BV510™ | V7 | Biolegend® |
| Il-4 | Intra | 11B11 | BV605™ | V10 | Biolegend® |
| Il-6 | Intra | MP5-20F3 | PerCP-eFluor™ 710 | B10 | Invitrogen™; eBioscience™ |
| Il-10 | Intra | JES5-16E3 | BV711™ | V13 | BD Biosciences© |
| Il-13 | Intra | 13A | PE-Cy5.5 | YG7 | Novus Biologicals© |
| Il-17 | Intra | TC11-18H10 | PerCP-Cy5.5 | B9 | BD Biosciences© |
| Tnf | Intra | MP6-XT22 | Pacific Blue (PB)™ | V3 | Biolegend® |
| Ifny | Intra | XMG1.2 | PE-Cy5 | YG5 | Abcam© |
| Perforin | Intra | CE2.10 | DyLight™ 405 | V2 | Novus Biologicals© |
| Granzyme b | Intra | QA16A02 | AF700™ | R4 | Biolegend® |

| | | | | | |
|---|---|---|---|---|---|
| Gm-csf | Intra | MP1-22E9 | APC | R1 | Biolegend® |

| Table S4. Immune Subset Phenotype Definitions | | |
|---|---|---|
| **Immune Subset** | **Phenotype (Gated on Single Live Cells)** | **References for Subset Identification (See Reference List Below Table)** |
| Bulk T cells | $Cd3^+$ | 1,2 |
| Naïve-Like Bulk T cells | $Cd3^+$ $Cd62l^+$ $Cd45ra^+$ | 3–10 |
| $T_{CM}$ Bulk T cells | $Cd3^+$ $Cd62l^+$ $Cd45ra^-$ | 3–7 |
| $T_{EM}$ Bulk T cells | $Cd3^+$ $Cd62l^-$ $Cd45ra^-$ | 3–7 |
| $T_{EF}$ Bulk T cells | $Cd3^+$ $Cd62l^-$ $Cd45ra^+$ | 3–10 |
| $T_{SCM}$ Bulk T cells | $Cd3^+$ $Cd62l^+$ $Cd45ra^+$ $Cd183^+$ | 3–22 |
| Bulk $Cd8^+$ T cells | $Cd3^+$ $Cd8a^+$ | 1,2 |
| Naïve-Like Bulk $Cd8^+$ T cells | $Cd3^+$ $Cd8a^+$ $Cd62l^+$ $Cd45ra^+$ | 1–10 |
| $T_{CM}$ Bulk $Cd8^+$ T cells | $Cd3^+$ $Cd8a^+$ $Cd62l^+$ $Cd45ra^-$ | 1–7 |
| $T_{EM}$ Bulk $Cd8^+$ T cells | $Cd3^+$ $Cd8a^+$ $Cd62l^-$ $Cd45ra^-$ | 1–7 |
| $T_{EF}$ Bulk $Cd8^+$ T cells | $Cd3^+$ $Cd8a^+$ $Cd62l^-$ $Cd45ra^+$ | 1–10 |
| $T_{SCM}$ Bulk $Cd8^+$ T cells | $Cd3^+$ $Cd8^+$ $Cd62l^+$ $Cd45ra^+$ $Cd183^+$ | 1–22 |
| Bulk $Cd4^+$ T cells | $Cd3^+$ $Cd4^+$ | 1,2 |
| Naïve-Like Bulk $Cd4^+$ T cells | $Cd3^+$ $Cd4^+$ $Cd62l^+$ $Cd45ra^+$ | 1–10 |

| | | |
|---|---|---|
| $T_{CM}$ Bulk $Cd4^+$ T cells | $Cd3^+$ $Cd4^+$ $Cd62l^+$ $Cd45ra^-$ | 1–7 |
| $T_{EM}$ Bulk $Cd4^+$ T cells | $Cd3^+$ $Cd4^+$ $Cd62l^-$ $Cd45ra^-$ | 1–7 |
| $T_{EF}$ Bulk $Cd4^+$ T cells | $Cd3^+$ $Cd4^+$ $Cd62l^-$ $Cd45ra^+$ | 1–10 |
| $T_{REG}$ | $Cd3^+$ $Cd4^+$ $Cd25^+$ $Cd127^-$ | 23–29 |
| Naïve-Like $T_{REG}$ | $Cd3^+$ $Cd4^+$ $Cd25^+$ $Cd127^-$ $Cd62l^+$ $Cd45ra^+$ | 3–10,23–29 |
| $T_{CM}$ $T_{REG}$ | $Cd3^+$ $Cd4^+$ $Cd25^+$ $Cd127^-$ $Cd62l^+$ $Cd45ra^-$ | 3–7,23–29 |
| $T_{EM}$ $T_{REG}$ | $Cd3^+$ $Cd4^+$ $Cd25^+$ $Cd127^-$ $Cd62l^-$ $Cd45ra^-$ | 3–7,23–29 |
| $T_{EF}$ $T_{REG}$ | $Cd3^+$ $Cd4^+$ $Cd25^+$ $Cd127^-$ $Cd62l^-$ $Cd45ra^+$ | 3–10,23–29 |
| $T_{CON}$ | $Cd3^+$ $Cd4^+$ $Cd25^{-/lo/+/hi}$ $Cd127^{+/hi}$ | 27–38 |
| Naïve-Like $T_{CON}$ | $Cd3^+$ $Cd4^+$ $Cd25^{-/lo/+/hi}$ $Cd127^{+/hi}$ $Cd62l^+$ $Cd45ra^+$ | 3–10,27–38 |
| $T_{CM}$ $T_{CON}$ | $Cd3^+$ $Cd4^+$ $Cd25^{-/lo/+/hi}$ $Cd127^{+/hi}$ $Cd62l^+$ $Cd45ra^-$ | 3–7,27–38 |
| $T_{EM}$ $T_{CON}$ | $Cd3^+$ $Cd4^+$ $Cd25^{-/lo/+/hi}$ $Cd127^{+/hi}$ $Cd62l^-$ $Cd45ra^-$ | 3–7,27–38 |

| | | |
|---|---|---|
| $T_{EF}$ $T_{CON}$ | $Cd3^{+}$ $Cd4^{+}$ $Cd25^{-/lo/+/hi}$ $Cd127^{+/hi}$ $Cd62l^{-}$ $Cd45ra^{+}$ | 3–10,27–38 |
| $T_{H1}$ | $Cd3^{+}$ $Cd4^{+}$ $Cd25^{-/lo/+/hi}$ $Cd127^{+/hi}$ $Cd183^{+}$ | 11–14,27–29,36–41 |
| Naïve-Like $T_{H1}$ | $Cd3^{+}$ $Cd4^{+}$ $Cd25^{-/lo/+/hi}$ $Cd127^{+/hi}$ $Cd183^{+}$ $Cd62l^{+}$ $Cd45ra^{+}$ | 3–14,27–29,36–41 |
| $T_{CM}$ $T_{H1}$ | $Cd3^{+}$ $Cd4^{+}$ $Cd25^{-/lo/+/hi}$ $Cd127^{+/hi}$ $Cd183^{+}$ $Cd62l^{+}$ $Cd45ra^{-}$ | 3–7,11–14,27–29,36–41 |
| $T_{EM}$ $T_{H1}$ | $Cd3^{+}$ $Cd4^{+}$ $Cd25^{-/lo/+/hi}$ $Cd127^{+/hi}$ $Cd183^{+}$ $Cd62l^{-}$ $Cd45ra^{-}$ | 3–7,11–14,27–29,36–41 |
| $T_{EF}$ $T_{H1}$ | $Cd3^{+}$ $Cd4^{+}$ $Cd25^{-/lo/+/hi}$ $Cd127^{+/hi}$ $Cd183^{+}$ $Cd62l^{-}$ $Cd45ra^{+}$ | 3–14,27–29,36–41 |
| $T_{H2}$ | $Cd3^{+}$ $Cd4^{+}$ $Cd25^{-/lo/+/hi}$ $Cd127^{+/hi}$ $Cd194^{+}$ | 27–29,36–41 |
| Naïve-Like $T_{H2}$ | $Cd3^{+}$ $Cd4^{+}$ $Cd25^{-/lo/+/hi}$ $Cd127^{+/hi}$ $Cd194^{+}$ $Cd62l^{+}$ $Cd45ra^{+}$ | 3–10,27–29,36–41 |

| | | |
|---|---|---|
| $T_{CM}$ $T_{H2}$ | $Cd3^{+}$ $Cd4^{+}$ $Cd25^{-/lo/+/hi}$ $Cd127^{+/hi}$ $Cd194^{+}$ $Cd62l^{+}$ $Cd45ra^{-}$ | 3–7,27–29,36–41 |
| $T_{EM}$ $T_{H2}$ | $Cd3^{+}$ $Cd4^{+}$ $Cd25^{-/lo/+/hi}$ $Cd127^{+/hi}$ $Cd194^{+}$ $Cd62l^{-}$ $Cd45ra^{-}$ | 3–7,27–29,36–41 |
| $T_{EF}$ $T_{H2}$ | $Cd3^{+}$ $Cd4^{+}$ $Cd25^{-/lo/+/hi}$ $Cd127^{+/hi}$ $Cd194^{+}$ $Cd62l^{-}$ $Cd45ra^{+}$ | 3–10,27–29,36–41 |
| $T_{H17}$ | $Cd3^{+}$ $Cd4^{+}$ $Cd25^{-/lo/+/hi}$ $Cd127^{+/hi}$ $Cd194^{+}$ $Cd196^{+}$ | 27–29,36–39,42–44 |
| Naïve-Like $T_{H17}$ | $Cd3^{+}$ $Cd4^{+}$ $Cd25^{-/lo/+/hi}$ $Cd127^{+/hi}$ $Cd194^{+}$ $Cd196^{+}$ $Cd62l^{+}$ $Cd45ra^{+}$ | 3–10,27–29,36–39,42–44 |
| $T_{CM}$ $T_{H17}$ | $Cd3^{+}$ $Cd4^{+}$ $Cd25^{-/lo/+/hi}$ $Cd127^{+/hi}$ $Cd194^{+}$ $Cd196^{+}$ $Cd62l^{+}$ $Cd45ra^{-}$ | 3–7,27–29,36–39,42–44 |
| $T_{EM}$ $T_{H17}$ | $Cd3^{+}$ $Cd4^{+}$ $Cd25^{-/lo/+/hi}$ $Cd127^{+/hi}$ $Cd194^{+}$ $Cd196^{+}$ $Cd62l^{-}$ $Cd45ra^{-}$ | 3–7,27–29,36–39,42–44 |

| | | |
|---|---|---|
| $T_{EF}$ $T_{H17}$ | $Cd3^+$ $Cd4^+$ $Cd25^{-/lo/+/hi}$ $Cd127^{+/hi}$ $Cd194^+$ $Cd196^+$ $Cd62l^-$ $Cd45ra^+$ | 3–10,27–29,36–39,42–44 |
| $T_{FH}$ | $Cd3^+$ $Cd4^+$ $Cd25^{-/lo/+/hi}$ $Cd127^{+/hi}$ $Cd185^+$ Pd-$1^+$ | 27–29,36–38,45,46 |
| Naïve-Like $T_{FH}$ | $Cd3^+$ $Cd4^+$ $Cd25^{-/lo/+/hi}$ $Cd127^{+/hi}$ $Cd185^+$ Pd-$1^+$ $Cd62l^+$ $Cd45ra^+$ | 3–10,27–29,36–38,45,46 |
| $T_{CM}$ $T_{FH}$ | $Cd3^+$ $Cd4^+$ $Cd25^{-/lo/+/hi}$ $Cd127^{+/hi}$ $Cd185^+$ Pd-$1^+$ $Cd62l^+$ $Cd45ra^-$ | 3–7,27–29,36–38,45,46 |
| $T_{EM}$ $T_{FH}$ | $Cd3^+$ $Cd4^+$ $Cd25^{-/lo/+/hi}$ $Cd127^{+/hi}$ $Cd185^+$ Pd-$1^+$ $Cd62l^-$ $Cd45ra^-$ | 3–7,27–29,36–38,45,46 |
| $T_{EF}$ $T_{FH}$ | $Cd3^+$ $Cd4^+$ $Cd25^{-/lo/+/hi}$ $Cd127^{+/hi}$ $Cd185^+$ Pd-$1^+$ $Cd62l^-$ $Cd45ra^+$ | 3–10,27–29,36–38,45,46 |
| NK cells | $Cd3^-$ $Nkp46^+$ | 47–50 |
| $S100b^+$ Only Astrocytes/Immune Cell | $Cd3^-$ $Gfap^-$ $S100b^+$ | 51–55 |

| | | |
|---|---|---|
| $Gfap^{+}$ Only<br>Astrocytes/Immune Cell | $Cd3^{-}$ $Gfap^{+}$ $S100b^{-}$ | 51–55 |
| $Gfap^{+}$ $S100b^{+}$<br>Astrocytes/Immune Cell | $Cd3^{-}$ $Gfap^{+}$ $S100b^{+}$ | 51–55 |
| DCs | $Cd3^{-}$ $Nkp46^{-}$ $Cd11c^{+}$ | 56–58 |
| Bulk $Cd11b^{+}$ Phagocytes | $Cd3^{-}$ $Nkp46^{-}$ $Cd11c^{-}$ $Cd11b^{+}$ | 59–62 |
| "Highly Active"<br>Microglia/Phagocytes | $Cd3^{-}$ $Nkp46^{-}$ $Cd11c^{-}$ $Cd11b^{+}$<br>$Iba1^{+}$ $Tmem119^{-}$ | 63–71 |
| "Active"<br>Microglia/Phagocytes | $Cd3^{-}$ $Nkp46^{-}$ $Cd11c^{-}$ $Cd11b^{+}$<br>$Iba1^{+}$ $Tmem119^{+}$ | 63–71 |
| "Steady State/Ramified"<br>Microglia/Phagocytes | $Cd3^{-}$ $Nkp46^{-}$ $Cd11c^{-}Cd11b^{+}$<br>$Iba1^{-}$ $Tmem119^{+}$ | 63–71 |
| Monocytes | $Cd3^{-}$ $Nkp46^{-}$ $Cd11c^{-}$ $Cd11b^{+}$<br>$Cd14^{+}$ $Cd68^{-/lo}$ | 59–62,72–76 |
| M1 Monocytes | $Cd3^{-}$ $Nkp46^{-}$ $Cd11c^{-}$ $Cd11b^{+}$<br>$Cd14^{+}$ $Cd68^{-/lo}$ $Cd86^{+}$ $Cd163^{-}$ | 59–62,72–76 |
| M2 Monocytes | $Cd3^{-}$ $Nkp46^{-}$ $Cd11c^{-}$ $Cd11b^{+}$<br>$Cd14^{+}$ $Cd68^{-/lo}$ $Cd86^{-}$ $Cd163^{+}$ | 59–62,72–76 |
| Macrophages | $Cd3^{-}$ $Nkp46^{-}$ $Cd11c^{-}$ $Cd11b^{+}$<br>$Cd14^{+}$ $Cd68^{+/hi}$ | 59,61,62,72,77 |

| | | |
|---|---|---|
| M1 Macrophages | Cd3$^{-}$ Nkp46$^{-}$ Cd11c$^{-}$ Cd11b$^{+}$ Cd14$^{+}$ Cd68$^{+/hi}$ Cd86$^{+}$ Cd163$^{-}$ | 59,61,62,73,77,78 |
| M2 Macrophages | Cd3$^{-}$ Nkp46$^{-}$ Cd11c$^{-}$ Cd11b$^{+}$ Cd14$^{+}$ Cd68$^{+/hi}$ Cd86$^{-}$ Cd163$^{+}$ | 59,61,62,73,77,78 |
| B cells | Cd3$^{-}$ Nkp46$^{-}$ Cd11c$^{-}$ Cd11b$^{-}$ Cd22$^{+}$ | 79–82 |

| Table S5. tSNE Analysis Details<br>Brain tSNE Analysis | | | | | |
|---|---|---|---|---|---|
| Cluster | WT Periphery Mean % within Each Mouse (N=5) with STD | GKI Periphery Mean % within Each Mouse (N=5) with STD | P-value to WT* | LKO Periphery Mean % within Each Mouse (N=5) with STD | P-value to WT* |
| Cd4+ Cd8+ T cells (Cd86+ Cd185+ Cd183+ Cd25+ Cd45ra+ Cd127+ Cd68+) | 0.36 ± 0.22 | 0.04 ± 0.05 | **0.0486** | 0.21 ± 0.25 | >0.9999 |
| Cd4+ T cells (Cd25+ Cd45ra+) | 0.07 ± 0.06 | 0.33 ± 0.21 | 0.3107 | 0.13 ± 0.26 | >0.9999 |
| Cd8+ T cells (Cd86+ Cd11c+ Cd183+ Cd25+ Cd45ra+ Cd127+ Cd68+) | 0.02 ± 0.02 | 0.64 ± 0.44 | **0.0179** | 0.01 ± 0.01 | 0.5433 |
| Cd8+ T cells (Cd86+ Cd25+ Cd45ra+ Cd68+) | 2.26 ± 0.82 | 6.31 ± 2.70 | 0.1017 | 1.12 ± 1.04 | 0.8665 |
| Cd8+ T cells (Cd86+ Cd183+ Cd25+ Cd45ra+ Cd127+ Cd68+) | 1.32 ± 0.69 | 4.59 ± 2.37 | **0.0294** | 0.37 ± 0.35 | 0.0873 |

| | | | | | |
|---|---|---|---|---|---|
| Cd8+ T cells (Cd86+ Cd185+ Cd11c+ Cd183+ Cd25+ Cd45ra+ Cd127+ Cd68+) | 0.36 ± 0.22 | 2.18 ± 2.29 | >0.9999 | 0.14 ± 0.22 | 0.5373 |
| Cd8+ T cells (Cd86+ Cd185+ Cd183+ Cd25+ Cd45ra+ Cd127+ Cd68+) | 1.32 ± 0.74 | 2.28 ± 1.09 | 0.8639 | 0.72 ± 1.34 | 0.7274 |
| Cd11b+ Tmem119+ Microglia (Cd62L+ Cd127+ Cd22+ Pd-1+) | 0.05 ± 0.04 | 0.02 ± 0.01 | 0.3718 | 0.04 ± 0.02 | 0.7958 |
| Cd11b+ Tmem119+ Microglia (Cd62l+ Cd183+ Cd68+) | 0.13 ± 0.02 | 0.09 ± 0.04 | 0.2494 | 0.35 ± 0.24 | 0.2760 |
| Cd11b+ Tmem119+ Microglia (Cd68+) | 0.41 ± 0.07 | 0.41 ± 0.22 | >0.9999 | 0.79 ± 0.10 | **0.0003** |
| DCs (Cd14+) | 0.10 ± 0.04 | 0.05 ± 0.04 | 0.2476 | 0.04 ± 0.03 | 0.0697 |
| Gfap+ Astrocytes | 0.06 ± 0.04 | 0.06 ± 0.05 | 0.9947 | 0.12 ± 0.05 | 0.1983 |
| **Periphery tSNE Analysis** | | | | | |

| Cluster | WT Periphery Mean % within Each Mouse (N=5) with STD | GKI Periphery Mean % within Each Mouse (N=5) with STD | P-value to WT* | LKO Periphery Mean % within Each Mouse (N=5) with STD | P-value to WT* |
|---|---|---|---|---|---|
| B Cells (Cd45ra+) | 52.95 ± 18.86 | 51.12 ± 4.58 | 0.9946 | 48.34 ± 4.51 | 0.9288 |
| Cd4+ Cd8+ T cells (Cd45ra+ Cd68+) | 0.20 ± 0.21 | 0.15 ± 0.05 | >0.9999 | 0.15 ± 0.15 | >0.9999 |
| Cd4+ T cells | 1.55 ± 2.09 | 3.68 ± 1.50 | 0.4127 | 3.39 ± 1.52 | 0.6093 |
| Cd4+ T cells (Pd-1+) | 3.32 ± 1.26 | 4.79 ± 1.97 | 0.4589 | 5.74 ± 1.13 | **0.0349** |
| Cd8+ T cells | 1.02 ± 1.00 | 2.00 ± 0.87 | 0.4127 | 2.16 ± 0.94 | 0.2691 |
| Cd8+ T cells (Cd11c+ Cd86+ Cd68+) | 0.10 ± 0.06 | 0.07 ± 0.04 | 0.7634 | 0.04 ± 0.03 | 0.2070 |
| Cd8+ T cells (Cd45ra+ Cd62l+) | 0.47 ± 0.24 | 0.46 ± 0.28 | >0.9999 | 0.85 ± 0.38 | 0.2461 |
| Cd8+ T cells (Cd45ra+ Cd68+) | 0.38 ± 0.08 | 0.27 ± 0.08 | 0.4127 | 0.21 ± 0.11 | **0.0327** |
| Cd8+ T cells (Cd45ra+) | 2.53 ± 1.07 | 3.06 ± 1.15 | 0.8303 | 4.75 ± 1.03 | **0.0281** |
| Cd8+ T cells (Cd68+) | 1.14 ± 0.29 | 1.01 ± 0.39 | 0.9896 | 1.09 ± 0.34 | 0.9063 |
| Cd11b+ Phagocytes (Cd62l+ Cd68+) | 1.29 ± 0.68 | 0.42 ± 0.14 | 0.2691 | 0.44 ± 0.13 | 0.1687 |
| Cd11b+ Phagocytes (Cd68+) | 5.63 ± 2.48 | 2.14 ± 0.32 | 0.0871 | 2.02 ± 0.94 | 0.0734 |
| Cd11b+ Phagocytes (Iba1+ Cd68+) | 1.36 ± 1.25 | 0.21 ± 0.16 | 0.2548 | 0.12 ± 0.07 | 0.2126 |

| | | | | | |
|---|---|---|---|---|---|
| DCs<br><br>(Cd68+) | 1.73 ± 0.36 | 2.44 ± 0.41 | 0.0643 | 2.56 ± 0.36 | 0.0532 |
| NK cells | 2.96 ± 1.41 | 3.65 ± 0.76 | 0.7212 | 2.01 ± 0.88 | 0.5364 |

* Parametric data: Brown-Forsythe and Welch ANOVA with a Dunnett's T3 multiple comparisons test

Non-parametric data: Kruskal-Wallis test with a Dunn's multiple comparison test
